# DNA Strand Breaks and Gaps Target Retroviral Binding and Integration

**DOI:** 10.1101/2021.11.17.469012

**Authors:** Gayan Senavirathne, Anne Gardner, James London, Richard Fishel, Kristine E. Yoder

**Affiliations:** Molecular Carcinogenesis and Chemoprevention Program, The James Comprehensive Cancer Center and Ohio State University, Columbus, OH 43210, US

## Abstract

Integration into a host genome is essential for retrovirus infection and is catalyzed by a nucleoprotein complex (Intasome) containing the viral integrase (IN) and reverse transcribed (RT) copy DNA (cDNA). Previous studies demonstrated DNA site recognition limited intasome integration. Using single molecule Förster resonance energy transfer (smFRET), we show Prototype Foamy Virus (PFV) intasomes pause at DNA strand breaks and gaps. The break/gap discontinuities are similar to base excision repair (BER) lesion-processing intermediates, which affect retrovirus integration *in vivo*. Pausing targeted site-directed integration at the break/gap without inducing intasome conformational alterations. An 8-oxo-guanine lesion normally processes by BER and a G/T mismatch or a +T nucleotide insertion that induce flexibility or a bend in the DNA backbone did not promote intasome pausing or targeted integration. These results suggest that repair intermediates can modulate dynamic intasome-DNA interactions which target retroviral integration.

---

Retroviruses must integrate into a host genome to cause clinically significant diseases such as AIDS or leukemia ^1^. Integration is catalyzed by a poorly understood retroviral pre-integration complex (PICs) that at a minimum consists of viral integrase (IN) multimers in complex with the two long terminal repeat (LTR) sequences that flank the viral cDNA, termed an intasome ^2-4^. The IN protein initially excises two nucleotides (nt) from the LTR ^5^ producing recessed 3’-OH’s that provide the substrate for integration. Consecutive IN-catalyzed S_N_2 strand transfer reactions then covalently link the recessed 3’-LTR ends across one major groove of a target DNA separated by 4-6 bp, depending on the retrovirus genera ^4,6,7^. When unraveled the newly integrated provirus is flanked by 4-6 nt gaps containing 2 nt 5’-flaps that are restored to fully duplex DNA by the host DNA repair machinery ^8-10^.

Alignment of numerous integration sites suggests that most retroviruses prefer to integrate at degenerate DNA sequences that for HIV-1 often positions a guanine (G) adjacent to the strand transfer site (T c **G** * G/t T N A C * C/a g A; uppercase favored, lowercase disfavored, * strand transfer site) ^11,12^. A large-scale siRNA screen identified base excision repair (BER) components as the most frequent DNA repair genes that altered HIV integration efficiency ^8^. Of these, mutation of oxidative damage BER glycosylases *OGG1* or *MUTYH* reduced HIV-1 integration as well as the preference for G and/or disfavored C adjacent to the strand transfer site ^9^. The OGG1 glycosylase principally excises 8-oxo-guanine (8-oxo-G) residues in DNA that are the most frequent oxidative damage in cells ^13,14^. The MUTYH glycosylase excises adenine (A) residues that are frequently misincorporated across from an 8-oxo-G lesion during replication ^15^. Mutation of the BER polymerase *β* (*Polβ*) also significantly decreased HIV-1 integration ^16^. However, the phenotype was linked to the 5’-deoxyribose phosphate (5’-dRP) lyase activity and not the polymerase activity of Pol*β* ^16^. The 5’-dRP lyase removes the deoxyribose sugar and associated phosphate leaving a 1 nt gap in the DNA after base excision by a glycosylase, ^17-19^. Interestingly, OGG1 also contains an intrinsic 5’-dRP lyase, while other glycosylases generally utilize the abasic endonuclease APE to introduce the strand break required to complete BER ^15^.

Historical studies have suggested that bent or flexible DNA is a favored target for retroviral integration *in vivo* ^20,21^ and *in vitro* ^22-29^. Prototype foamy virus (PFV) is a member of the *spumavirus* genera and has been extensively studied since it shares similar catalytic site geometry, chemistry and therapeutic inhibitor sensitivity with the pathogenic lentivirus HIV-1 ^30-32^. Structures of the PFV intasome target capture complex (TCC) and the strand transfer complex (STC) have been solved and represent key steps that provide a biophysical window into retroviral integration ^22,30^. Real-time single molecule imaging has detailed the dynamic progressions between PFV intasome and target DNA ^23,29^. These studies concluded that DNA site recognition limited PFV integration ^23^. However, the DNA configurations that drive integration site choice remain enigmatic.

## RESULTS

### PFV intasomes pause at breaks and gaps within target DNA

A single molecule Förster resonance energy transfer (smFRET) system ^33^ was developed to probe the real-time interactions between a purified PFV intasome with DNA containing a variety of DNA lesions and BER intermediates that introduce localized bends or flexibility into DNA ^34-37^ (**Fig. 1a**; **Extended Data Table 1**). PFV intasomes were assembled with pre-processed recessed 3’-OH viral 5’-LTR oligonucleotides (vDNA) containing a Cy3 FRET-donor flourophore on the non-transferred strand 11 bp from the 3’-end (Cy3-PFV). We confirmed that concerted integration of the Cy3-PFV intasomes into a supercoiled target DNA mimicked unlabeled PFV intasomes, suggesting that the Cy3-labeled vDNA does not affect integration activity (**Extended Data Fig. 1**) ^23^.

**Figure 1.**
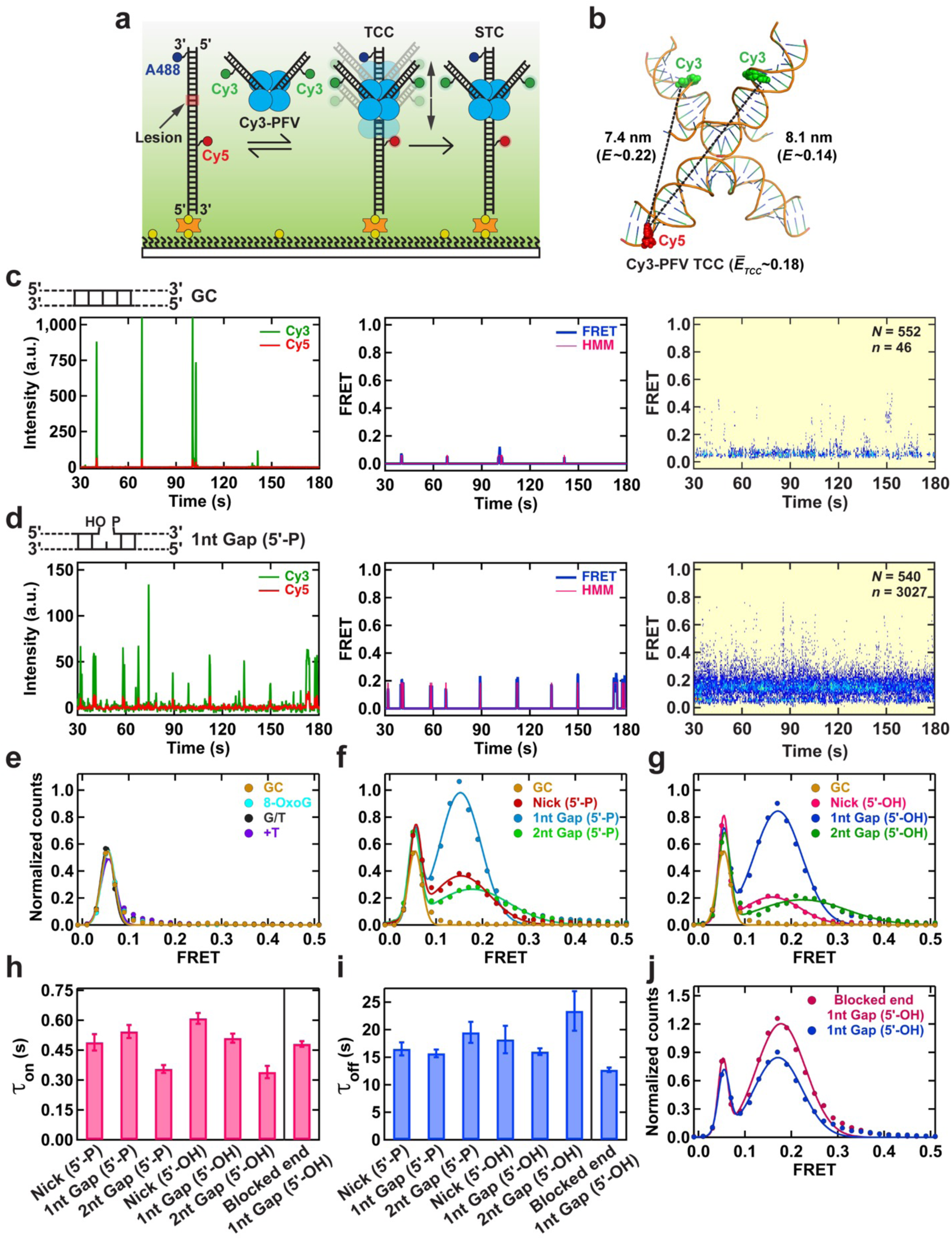
Real-Time PFV Intasome Target Capture Dynamics. (**a**) An illustration of the smFRET experimental setup for visualizing the target capture by a PFV intasome labeled with Cy3 at non-transfer strands (Cy3-PFV). The intasome was introduced in real-time onto DNA targets containing Alexa Fluor 488 (A488) and Cy5. (**b**) Positioning of the DNAs within the PFV TCC structure (PDB: 3OS2) showing the fluorophore positions on the vDNAs (Cy3) and the target DNA (Cy5), respectively. The estimated inter-dye distances, corresponding FRET efficiencies, and the average FRET value (*E*_*TCC*_) of a mixture of molecules containing a single Cy3 on the left or right vDNA. (**c**,**d**) A representative intensity trajectory (left), the corresponding FRET trajectory with the Hidden Markov Model (HMM) fit (middle), and the post-synchronized histogram (PSH) (right) displaying intasome interactions with GC duplex target DNA (**c**), and target DNA containing a 1nt Gap (5’-P) (**d**). Inset shows the total number of DNA molecules (*N*) analyzed for each substrate and the total number of transitions with >0.1 FRET (*n*). (**e**) Normalized smFRET histograms and Gaussian fits (**Extended Data Table 3 and 4**) showing the distributions of FRET efficiency for target DNAs containing a GC duplex, 8-Oxo-G lesion, G/T mismatch, +T insertion. (**f**,**g**) Normalized smFRET histograms and Gaussian fits (**Extended Data Table 3 and 4**) showing the distributions of FRET efficiency for target DNAs containing single strand scission (Nick), a 1 nt Gap, and a 2 nt Gap with a 5’-phosphate (5’-P) (**f**) or 5’-hydroxyl (5’-OH) (**g**). The GC duplex target DNA was included for comparison. (**h**,**i**) The TCC binding lifetime (*τ*_on_) (**h**) and lifetime of the dissociated state (*τ*_off_) (**i**) for different target DNA substrates. Error bars represent the errors from the fittings of dwell time distributions (**Extended Data Fig. 8)**. (**j**) Normalized smFRET histograms and their Gaussian fits (**Extended Data Table 3 and 4**) showing the FRET distribution for the 1nt Gap (5’-OH) DNA with or without blocking the open DNA end.

A 60 bp target DNA containing defined lesions was synthesized that contained a Cy5 FRET-acceptor flourophore located on the undamaged strand 11 bp 5’ of the lesion as well as an Alexa488 marker flourophore on the damaged strand 3 bp from the 3’ end (**Fig. 1a, Extended Data Table 1**). This target DNA was bound to a passivated smFRET flow cell surface at the 5’-end of the lesion-containing strand and purified Cy3-PFV intasomes infused and illuminated with a 532 nm laser. Binding of the Cy3-PFV intasomes near the lesion results in Cy5 (660 nm) FRET emission events consistent with dynamic TCC interactions (**Fig. 1b and 1c**, left; 100 msec frame rate). Background correction and Hidden Markov Modeling (HMM) generated a transition density plot ^38^ that was used to determine FRET efficiency (*E*) and reversibility (**Fig. 1c**, middle; **Extended Data Fig. 2 and 3**). The total number of FRET events (n) from numerous DNA molecules (N) were aligned to the PFV intasome infusion time to produce a post-synchronized histogram (N = 552, n = 46; **Fig. 1c**, right) ^39^. A narrow range of Cy5 fluorescence (*E*_*pseudo*_ ∼ 0.06) was detected in both the transition density plot and post-synchronized histogram that after inspection of the real-time movies were found to represent excursions of free Cy3-PFV intasome aggregates across the evanescent field containing the target DNA (**Extended Data Movie 1**). These events were easily recognized since they saturated the Cy3 channel and bled into the Cy5 channel producing a pseudo-FRET signal (**Fig. 1c**, right). Rare higher FRET events (*E* > 0.1) were detected that appeared to correspond to PFV intasome binding and dissociation events near the Cy5 fluorophore on the DNA (**Fig. 1c**, right). A similar pattern was observed when an 8-oxo-G/C, a G/T mismatch or +T nucleotide insertion was present in the duplex target DNA (**Extended Data 4a-c**). The lack of significant FRET events above the pseudo-FRET background suggests very little if any specific PFV intasome binding to these target DNA substrates.

In contrast, a DNA substrate containing a 1 nt gap with a 3’-hydoxyl and 5’-phosphate [1 nt GAP (5’-P)] similar to a 5’-dRP lyase BER intermediate consistently displayed bursts of Cy3-PFV binding events accompanied by Cy5 FRET events (**Fig. 1d**, left). Inspection of real-time movies confirmed that these FRET events represented PFV intasome-lesion interactions arising from colocalized diffraction-limited Cy3-Cy5 spots (**Extended Data Movie 2**). The FRET bursts from individual traces (**Fig. 1d**, center) converged into a distinct ensemble FRET peak that was clearly separate from the pseudo-FRET background (*E*_*1nt Gap (5’-P)*_ ∼ 0.16 ± 0.05; N = 540, n = 3267; **Fig. 1d**, right; **Extended Data Table 4**). These events are consistent with the formation of a stably but transient TCC, with the PFV intasome bound near the DNA lesion (**Fig. 1b**; **Fig. 1d**). DNA substrates containing a single-strand scission [Nick (5’-P); *E*_*Nick (5’-P)*_ ∼ 0.15 ± 0.08; N = 549, n = 1084], a 2 nt gap [2 nt GAP (5’-P); *E*_*2nt Gap(5’-P)*_ ∼ 0.18 ± 0.10; N = 474, n = 1175] as well as a 5’-phosphate free nick (*E*_*Nick (5’-OH)*_ ∼ 0.17 ± 0.07; N = 558, n = 507), 1 nt gap (*E*_*1nt Gap (5’-OH)*_ ∼ 0.19 ± 0.05; N = 496, n = 2784) and 2 nt gap (*E*_*2nt Gap (5’-OH)*_ ∼ 0.23 ± 0.11; N = 540, n = 799) also yielded significant FRET events consistent with the formation of a TCC (**Extended Data Fig. 4d-h**; **Extended Data Table 2**). Importantly, the number of FRET transitions per molecule also increased with DNA substrates that displayed clear binding events (**Extended Data Fig. 5**). The consistent peak of FRET efficiency suggests the Cy3-PFV intasomes bind within a narrow distance around the DNA lesions.

To quantify the relative binding efficiency of PFV intasomes to the DNA lesions, we normalized the frequency of TCC FRET binding events to the pseudo-FRET excursion events that appear relatively constant between target DNA substrates at a fixed Cy3-PFV intasome concentration and frame rate (**Fig. 1e-g**; **Extended Table 3**). As expected, there were few FRET events outside the normalized pseudo-FRET peak when duplex DNA, DNA containing an 8-oxo-G/C lesion, a G/T mismatch or a +T nucleotide insertion were examined (*E* > 0.1; **Fig. 1e**; **Extended Data Fig. 6a-d**). Since the G/T mismatch introduces significant local flexibility into the DNA ^40,41^ and the +T insertion produces a relatively stable 22° DNA bend ^42^, these observations suggest that the preference for flexible or bent DNA by retrovirus intasomes ^20,43,44^ may be more subtle than previously appreciated.

In contrast, a nick, 1 nt gap or 2 nt gap, with or without the 5’-P, displayed normalized FRET peaks that were clearly separated from the pseudo-FRET background (**Fig. 1f,g**; **Extended Data Fig. 6e-j**). The relative frequency of these TCC events suggested that Cy3-PFV preferred a 1 nt Gap >> nick ≥ 2 nt Gap. We noted a broadening of the normalized TCC FRET peak as well as an increased average FRET efficiency with target DNA containing a 2 nt GAP (**Fig. 1f,g**). Because the 2 nt Gap extends further toward the Cy5 FRET acceptor, these results suggest an increase in the number of different asymmetric binding events that on average bind closer to the Cy5 fluorophore and 5’ of the gap. The post-synchronized histograms and the relative binding efficiency pattern of TCC FRET events was similar at a 1 sec frame rate (**Extended Data Fig. 7**; **Extended Table 4**). Because of the potential for enhanced binding resolution, we chose a 100 msec frame rate for the majority of these studies.

Using HMM we determine the binding (*τ*_on_) and dissociation (*τ*_off_) kinetics (**Extended Data Fig. 8a**). As expected, the binding events to an 8-oxo-G/C, a G/T mismatch or a +T nucleotide insertion were extremely rare, often < 1 event per molecule over the 3 min observation window, making any *τ*_on_ and *τ*_off_ values statistically insignificant (**Extended Data Fig. 5a-d**; **Extended Data Fig. 8b**). However, the 1 nt GAP (5’-P) and 1 nt Gap (5’-OH) experienced an average of ∼6 binding events per molecule with a distribution of 1-14 binding events for the vast majority of molecules (**Extended Data Fig. 5f**,**i**). Less, but statistically significant binding events were recorded for DNAs containing a nick and 2 nt gap regardless of the presence of a 5’-phosphate within the lesion (**Extended Data Fig. 6e**,**g**,**i**,**j**). Fitting these events to a single exponential decay resulted in a *τ*_on_ that varied from 0.3 - 0.75 sec and an *τ*_off_ that varied from 7 - 24 sec (**Fig. 1h,i**; **Extended Data Fig. 8c-h**). We note that the *τ*_on_ events for the 2 nt gap substrates approached the frame rate which appears to reduce the number of recorded events, and in the case of the 2 nt GAP (3’-OH) artificially increased the *τ*_off_ (**Fig. 1h,i**; **Extended Data Fig. 8e,h**). Blocking the open end of the 1nt GAP (5’-OH) DNA substrate to trap intasomes diffusing along the DNA ^45^ did not significantly alter the distribution of events per molecule or the post-synchronized FRET histogram (**Extended Data Fig. 9a,b**). However, while the *τ*_on_ remained largely constant the *τ*_off_ significantly decreased (**Fig. 1h.i**; **Extended Data Fig. 9c-e**), resulting in an increased number of normalized lesion binding events (**Fig. 1j**). These observations suggest that at least a fraction of the Cy3-PFV intasomes that form a TCC may dissociate by sliding off the ends of the lesion containing oligonucleotide.

### PFV intasomes catalyze strand transfer at the target DNA pause site

Integration by the Cy3-PFV intasomes forms a strand transfer complex (STC) that results in prolonged FRET events (**Fig. 2a,b**). These sustained FRET events often ended with photobleaching of the Cy3 or Cy5 fluorophores on the intasome or the target DNA substrate (**Fig. 2c**). Post-synchronized histograms displayed well defined STC FRET events for several of the DNA substrates, where both the frequency and FRET efficiency could be quantified (**Fig. 2d,e**; **Extended Data Table 2**). We found that the FRET efficiency of TCC events (*E*_*TCC*_) was nearly identical to the STC events (*E*_*STC*_; **Extended Data Table 2**). Where it differed, for example the Nick (5’-P) DNA substrate, the number of events were extremely low (*n* < 5) making interpretation of the histograms impractical (**Extended Data Fig. 10)**. The consistency in FRET efficiencies is consistent with the conclusion that the TCC pausing events contribute to the formation of an STC product.

**Figure 2.**
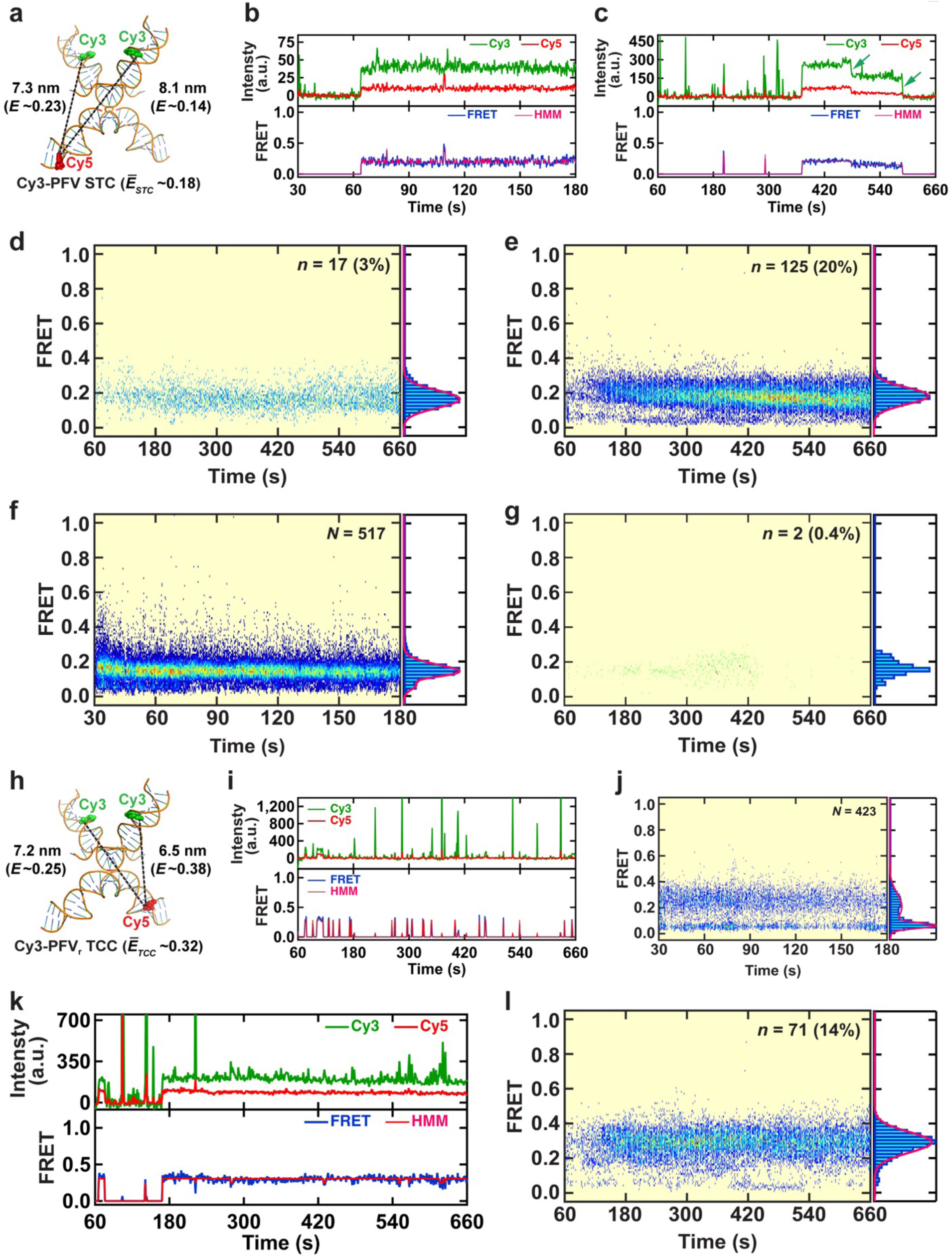
Real-Time PFV Intasome Catalyzed Strand Transfer. (**a**) Illustration of the DNA configuration found in the PFV strand transfer complex (STC) crystal structure (PDB: OS0) showing the fluorophore positions on the vDNAs (Cy3) and the target DNA (Cy5), respectively. The estimated inter-dye distances, corresponding FRET efficiencies, and the average FRET value (*E*_*STC*_) of a mixture of molecules containing a single Cy3 on the left or right vDNA. (**b,c**) Representative intensity trajectories and corresponding FRET trajectories with Hidden Markov Model (HMM) fits showing stable strand transfer by a Cy3-PFV intasome. Green arrows mark the photobleaching of the Cy3 FRET donor. (**d,e**) The post-synchronized histogram (PSH) and smFRET histograms corresponding to STC events for target DNAs containing a 1nt Gap (5’-P) (**d**) or 1nt Gap (5’-OH) (**e**). The Gaussian fits to the FRET histograms are shown as red lines. Data were collected at 100 msec frame rate (**b**) or 1 sec frame rate (**c-e**) resolution. Inset shows the total number of transitions with >0.1 FRET (*n*) and the percentage (%) of strand transfer events. (**f,g**) The post-synchronized histogram (PSH) and smFRET histograms corresponding to TCC (**f**) and STC (**g**) events for Cy3-PFV-ddA interacting with 1nt Gap (5’-OH) target DNA. Data were collected at 100 msec frame rate (**f**) and 1 sec frame rate (**g**) using a smFRET setup similar to **Fig. 1a**. Inset shows the total number of DNA molecules analyzed (*N*) (**f**); and the number of stable transitions with >0.1 FRET (*n*) and the percentage (%) of STC events. (**h**) Illustration of the DNA configuration found in the PFV strand transfer complex (STC) crystal structure (PDB: OS0) showing the fluorophore positions on the vDNAs (Cy3) and the reverse-Cy5 (R-Cy5) target DNA, respectively. The estimated inter-dye distances, corresponding FRET efficiencies, and the average FRET value (*E*_*TCC*_) of a mixture of molecules containing a single Cy3 on the left or right vDNA. (**i**) A representative intensity trajectory (top) and the corresponding FRET trajectory (bottom) with the Hidden Markov Model (HMM) fit showing Cy3-PFV binding to R-Cy5 target DNA containing a 1nt Gap (5’-OH). (**j**) The post-synchronized histogram (PSH, left) and the smFRET histogram (right) produced by averaging the total number (*N*) of TCC smFRET traces. (**k**) A representative intensity trajectory (top) and the corresponding FRET trajectory (bottom) with Hidden Markov Model (HMM) fit showing Cy3-PFV strand transfer into R-Cy5 target DNA containing a 1nt Gap (5’-OH). (**I**) The post-synchronized histogram (PSH, left) and the smFRET histogram (right) corresponding to the total number (*n*) and percentage (%) of STC events. The data in (**i-l**) were collected at 1 sec frame rate using a smFRET setup similar to the **Fig. 1a**.

To confirm that the STC events formed a covalent integration structure we assembled PFV intasomes with viral donor oligonucleotides containing a terminal di-deoxy-adenosine (ddA) (**Fig. 2f,g**; **Extended Data Table 5**). We used the1 nt Gap (5’-OH) target DNA as a model since it displayed the most frequent TCC and STC events. As expected, we observed significant numbers of TCC events that displayed a similar FRET efficiency to *wild type* PFV intasome interactions (compare **Fig. 2f** to **Extended Data Fig. 4g**; **Extended Data Table 2**). However, only two possible STC events were recorded on *N* = 564 DNA molecules (STC_1nt Gap (5’-OH)-ddA_ = 0.35 ± 0.2%) compared to 20% with *wild type* Cy3-PFV (**Fig. 2g**; **Extended Data Table 5**). This >50-fold reduction in STC events with Cy3-PFV-ddA intasomes strongly suggests that the prolonged FRET events represent covalent STC formation.

To determine whether the positioning of the fluorophore on the target DNA influenced PFV intasome interactions, we moved the Cy5 to 11 bp 3’ of the 1 nt Gap (5’-OH) (**Fig. 2h**; **Extended Data Table 1**). Similar TCC FRET events (**Fig. 2i,j**) and STC FRET events (**Fig. 2k,l**) were observed compared to the target DNA containing Cy5 located 11 bp 5’ of the 1 nt Gap (5’-OH) (**Extended Fig. 4g**; **Fig. 2e**). Intriguingly, the FRET efficiency of the TCC and STC events with the Cy5 located 11 bp 3’ of the 1 nt Gap (5’-OH) was greater than the FRET efficiency of the TCC and STC events with the Cy5 located 11 bp 5’ of the 1 nt Gap (5’-OH). These results are consistent with the conclusion that the Cy3-PFV intasome binds asymmetrically to the lesion-containing target DNA, with a preferred binding on the 3’-side of the 1 nt Gap (5’-OH). Nevertheless, the frequency of STC events appeared similar to the normalized counts of TCC FRET events (compare **Fig. 1e-g**, *E* > 0.1 with **Fig. 3a**, smFRET; **Extended Table 5**). These results suggest that the fluorophore location has little or no effect on TCC or STC events. Taken together we conclude that transient TCC events contribute to increased STC products.

**Figure 3.**
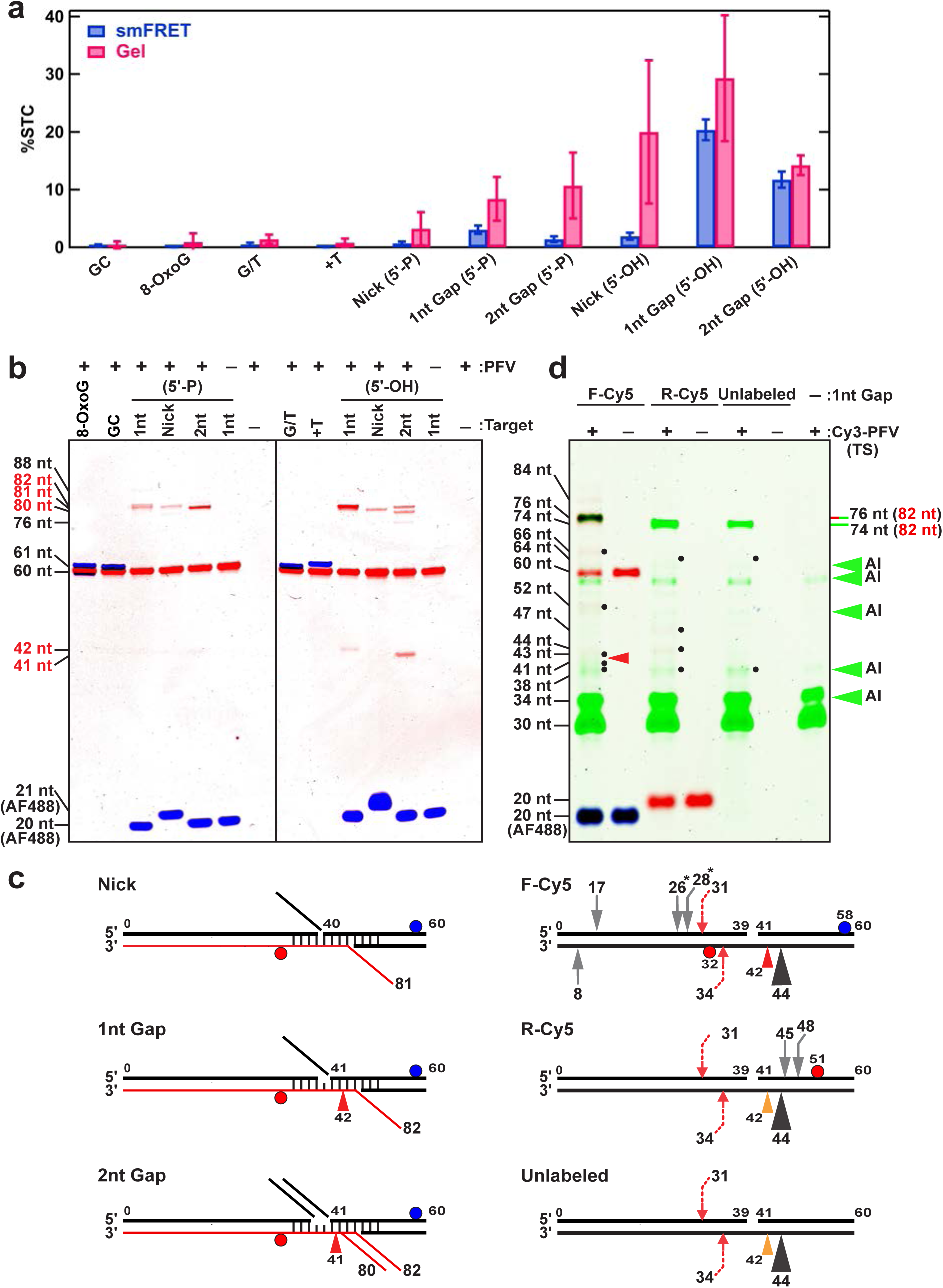
Analysis of PFV strand transfer activity. (**a**) Quantification of single-molecule and gel analysis of STC integration activity on different DNA targets (%). The smFRET data were generated using Cy3-PFV, and the gel data were obtained using unlabeled PFV with Cy5 and Alexa Fluor 488 labeled target DNAs. The error bars in the smFRET data reflect vn/N×100%, where *n* is the number of strand transfer events and *N* is the total number of DNA molecules analyzed. Error bars in the gel data are the standard deviations from triplicates. Differences in absolute frequency reflect different intasome reaction concentrations in smFRET (5 nM) or gel analysis (25 nM). (**b**) Representative denaturing PAGE gels from bulk integration studies. The lengths of ssDNAs derived from a Sanger sequencing ladder are shown. The blue and red bands show DNA fragments containing Alexa Fluor 488 (A488) and Cy5, respectively. The target DNA substrates used for each lane are shown above (**Extended Data Table 1**). (**c**) Schematics showing the major strand transfer event and predicted ssDNA length (red line), or aborted strand break (red arrowhead) exhibited by unlabeled PFV. (**d**) Gel analysis (top) and integration sites (bottom) of PFV intasomes labeled with Cy3 on the vDNA transferred strand [Cy3-PFV (TS)]. Integration into different 1nt Gap (5’-OH) substrates containing a forward Cy5 label (F-Cy5), reverse Cy5 label (R-Cy5 or unlabled are shown in Key on to and illustrated below. The cyan, green and red color gel bands correspond to DNA fragments containing Alexa Fluor 488 (AF488), Cy3, and/or Cy5. The brown color bands contain both Cy3 and Cy5. The black dots indicate the primary product resulting from strand integration and red arrowhead the 42 nt aborted strand transfer product (see: Panel b,c). Calculated integration sites are shown below for each target DNA. Large black arrow indicates major (>90%) half-site integration product similar to major bands in Panel b; red dashed arrows indicate minor concerted strand transfer product; gray arrows show minor half-site products. The numbers indicate the positions of integration relative to the 5’-end of the top target DNA strand. * Indicates products that have more than one integration site solution (see text).

### DNA breaks and gaps target site-specific PFV integration

To map the STC events we examined strand transfer products by denaturing gel electrophoresis (DGE; **Fig. 1a**, Gel; **Fig. 3b,c**). These studies incubated unlabeled PFV intasomes with the target DNAs utilized in the smFRET analysis (**Fig. 1a,b**; **Extended Data Table 1**). No integration events were observed in control reactions (-PFV) or with fully duplex target DNA (G/C) as well as target DNA containing an 8-Oxo-G (8-Oxo-G), G/T mismatch (G/T) or a +T insertion (+T). However, additional DNA bands of varying intensity were observed with a target DNA containing a Nick (5’-P or 5’-OH), a 1 nt Gap (5’-P or 5’-OH) or a 2 nt Gap (5’-P or 5’-OH) that are consistent with covalent PFV integration products (**Fig. 3b**). The pattern and frequency of STC events observed by smTIRF generally mimicked the Gel analysis (**Fig. 3a**), with differences in the absolute frequency of integration attributable to the lower concentration of PFV intasomes utilized in smFRET to moderate background fluorescence (5 nM smTIRF compared to 25 nM for Gel analysis).

Most strand transfer events observed by DGE were the result of half-site integration (**Fig. 3b**). For example, the target DNA containing a 1 nt Gap, Nick and 2 nt Gap with a 5’-P resulted in major Cy5-DNA bands of 82, 81, and 80 nt, respectively (**Fig. 3b**, left; see Key above). These correspond to covalent strand transfer of the 38 bp vDNA into the non-lesion containing Cy5-labeled strand 44, 43, 42 nt from the 3’-end, respectively (**Fig. 3c**). Since PFV integration events are separated by 4 bp ^22,30^, this strand transfer event would place the concerted strand transfer location on the lesion-containing strand at the 5’-side of the missing nucleotide within the 1 nt Gap, at the Nick or at the 5’-side of the second missing nucleotide of the 2 nt Gap. In all these cases there is no phosphate bond to complete the isoenergetic strand transfer chemistry. A similar integration pattern was detected with target DNAs containing a 1 nt Gap, Nick and 2 nt Gap with a 5’-OH (**Fig. 3b**, right). Importantly, the asymmetric location of these integration products corresponds with the asymmetric binding predicted by TCC and STC FRET events.

There were curious differences in band intensity and/or number when the 5’-P and 5’-OH lesions were compared (**Fig. 3b**). These include a substantial increase in the 82 nt product with the target DNA containing a 1 nt Gap (5’-OH) and an apparent splitting of intensity between 82 nt and 80 nt products with the target DNA containing a 2 nt Gap (5’-OH) (**Fig. 3b**, right). The 80 nt and 82 nt products with the 2 nt Gap (5’-OH) is consistent with ∼50% of the PFV intasomes positioning next to the 3’-OH and ∼50% positioning next to the 5’-side of the second missing nucleotide similar to the 2 nt Gap (5’-P) (**Fig. 3c**). These results appear internally consistent with the broadening and shift to increased FRET efficiency of STC events observed by smFRET with target DNA containing a 2 nt Gap (5’-OH), and suggest that the presence of a 5’-P can influence the location of half-site strand transfer events by altering the stable positioning of PFV intasomes.

In addition, a 42 nt and 41 nt DNA product was also detected with the 1 nt Gap (5’-OH) and 2 nt Gap (5’-OH), respectively (**Fig. 3b**, right; **Fig. 3c**). These DNA products appear consistent with aborted strand transfer events that leave a strand scission on the non-lesion containing Cy5-labeled strand. In both cases, we would be unable to detect a corresponding strand transfer on the fluorophore-deficient lesion-containing strand. Moreover, a weak but clearly visible DNA product of ∼76 nt with the target DNAs containing a 2 nt Gap (5’-OH) (**Fig. 3b**). This product could conceivable signify a concerted integration event since the second strand transfer should occur 35 nt from the 5’-end on the lesion-containing strand, which would also be undetectable with unlabeled PFV intasomes.

To determine whether additional integration events occur on the target DNA, we assembled PFV intasomes containing Cy3-labeled vDNA on the transferred strand (Cy3-PFV (TS); **Extended Data Table 1**). Cy3-PFV (TS) integration events transfer a 30 nt vDNA to an unlabeled target DNA (unlabeled), a target DNA containing a Cy5-fluorophore located 5’ of the lesion on the non-lesion containing strand (F-Cy5) or a target DNA containing a Cy5-fluorophore located 3’ on the lesion-containing strand (R-Cy5) (**Extended Data Table 1**). We examined the 1 nt Gap (5’-OH) and 2 nt Gap (5’-OH) target DNA substrates since they initially revealed integration products other than the major half-site strand transfer events (**Fig. 3b**). The principal Cy3-PFV (TS) integration events (>90%) produced fragments of 76 nt or 74 nt for both the 1 nt Gap (5’-OH) and 2 nt Gap (5’-OH) (**Fig. 3d**; **Extended Data Fig. 11a**). Subtracting the 30 nt vDNA from the strand transfer product and accounting for a 2 nt apparent size increase when both a Cy3 and Cy5 fluorophore are present when compared to Cy3-labeled molecular weight markers, the calculated integration events occurred at 44 nt on the non-lesion containing strand (**Fig. 3d**; **Extended Data Fig. 11a,b**; black arrowhead). This is the identical location that produced the 80 nt and 82 nt half-site integration products with the unlabeled PFV intasomes (**Fig. 3b**). We also detected background levels of the 41 nt and 42 nt aborted strand transfer product with the F-Cy5 target DNA substrates (**Fig. 3d**; **Extended Data Fig. 11a,b**; red arrowhead) that would be undetectable with the R-Cy5 and Unlabeled Target DNAs (**Fig. 3d**; **Extended Data Fig. 11**; orange arrowhead).

Minor Cy3-PFV (TS) integration products were identified with 1 nt Gap (5’-OH) and 2 nt Gap (5’-OH) target DNA substrates (**Fig. 1d**, black dots), that could be enhanced by increasing the image contrast (**Extended Data Fig. 11a,b**). These additional integration events accounted for < 5% of the target DNA integration products, with the majority localized around the lesion (gray arrows) and including a distinct concerted integration product adjacent to the lesion (red dashed arrow; **Fig. 1d**, bottom). We note that half-site integration products mapped to 26 nt and 28 nt on the F-Cy5 target DNA could also be localized to the non-lesion containing strand at 11 nt and 13 nt, respectively, and are therefore marked with a star (**Fig. 1d**, bottom). The 26, 28, 45 and 48 integration products appeared to be equidistant pairs from with the Cy5 fluorophore location (**Fig. 1d**), suggesting other DNA lesions might also enhance retroviral integration. More products could be visualized with increased image contrast but accounted for an even more minor fraction of the events. Together, these results confirm the conclusion that target DNA mimicking BER intermediates significantly enhances integration at the site of the lesion.

### Target DNA binding and integration does not alter the PFV intasome structure

Whether TCC and STC events provoke a conformational transition within the PFV intasome is unknown. To address this question, we designed a smFRET system in which alterations in the relative geometry of the vDNA could be monitored with high precision (**Fig. 4a**). PFV intasomes were assembled with two donor vDNAs, one containing a Cy3 fluorophore and the other containing a Cy5 fluorophore (Cy3/Cy5-PFV; **Fig. 4a,b**; **Online Methods**). Structural studies suggest that the position of the vDNA within the TCC and STC are nearly identical with similarly identical calculated FRET efficiency (**Fig. 4b**) ^22,30^. These structures have retained IN subunits that appear to localize the vDNA ^22,30^. The dynamic stability of these IN subunits during TCC and STC formation has not been determined ^22,30^.

**Figure 4.**
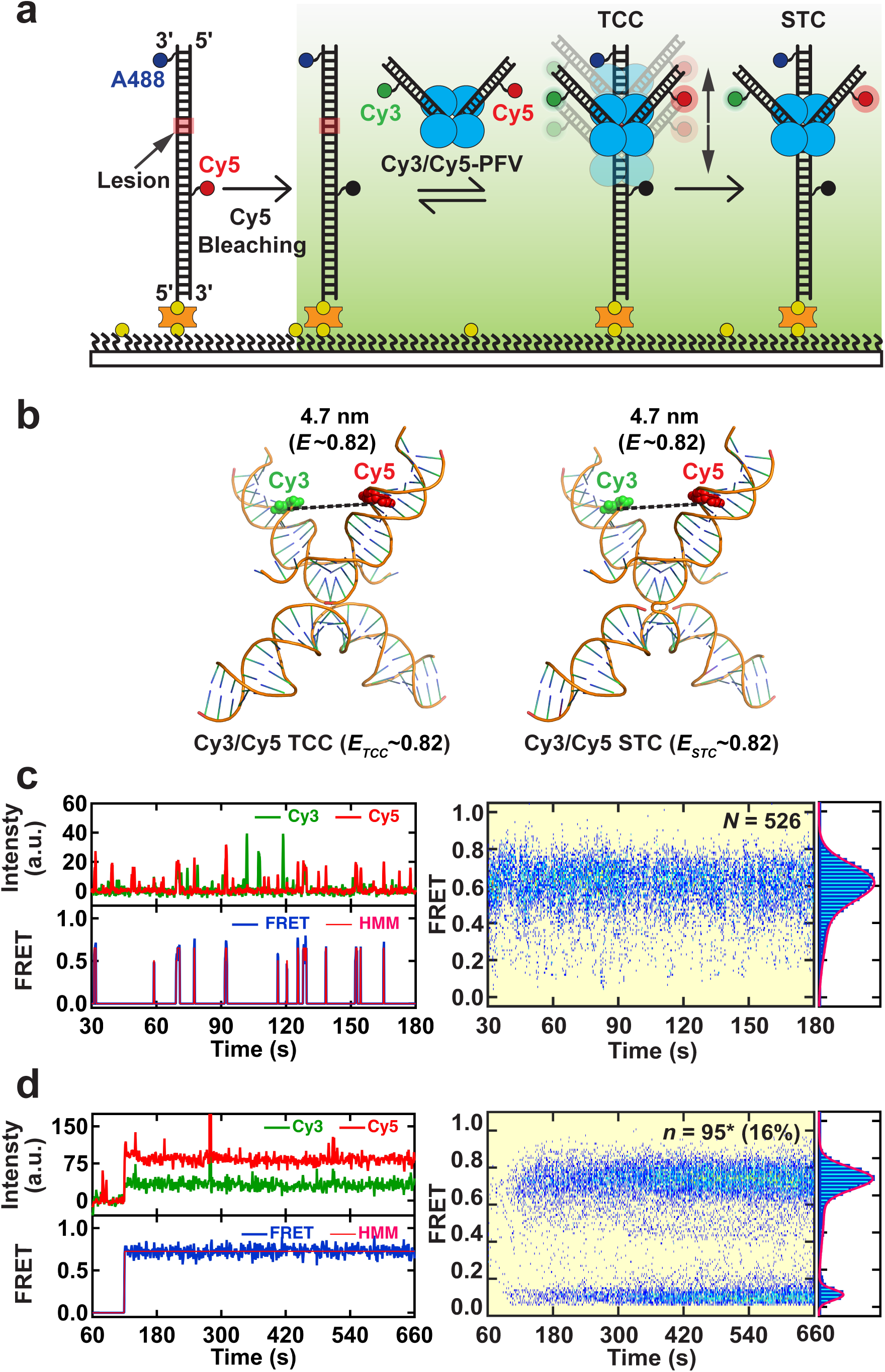
Probing the structural dynamics of PFV intasome during target capture and strand transfer. (**a**) An illustration of the smFRET experimental setup for for visualizing the structural dynamics of PFV intasome during TCC and STC formation. A PFV intasome labeled with Cy3 and Cy5 at the non-transfer strands (Cy3/Cy5 PFV) was introduced after photobleaching Cy5 on the substrates. (**b**) Illustration of the DNA configuration found in the PFV target capture complex (TCC) crystal structure (PDB: 3OS2) and strand transfer complex (STC) crystal structure (PDB: OS0) showing the fluorophore positions on the vDNAs (Cy3 and Cy5). The estimated inter-dye distances, corresponding FRET efficiencies, and the average FRET value (*E*_*TCC*_ and *E*_*STC*_) are shown. (**c**) Left: A representative TCC intensity trajectory (top) and the resulting FRET trajectory (bottom) with the Hidden Markov Model (HMM) fit showing Cy3/Cy5-PFV binding to a 1nt Gap (5’-OH) target DNA (left). Right: The post-synchronized histogram (PSH, left) and the smFRET histogram (right) generated by averaging the total number (*N*) of TCC FRET traces. (**d**) Left: A representative STC intensity trajectory (top) and the resulting FRET trajectory (bottom) with the Hidden Markov Model (HMM) fit showing Cy3/Cy5-PFV integration into a 1nt Gap (5’-OH) target DNA (left). Right: The post-synchronized histogram (PSH, left) and the smFRET histogram (right) generated by averaging the total number (*n*) of STC Cy3/Cy5-PFV FRET traces (* Cy3/Cy3-PFV events are included in the total number of STC traces). The percentage (%) is the efficiency of Cy3/Cy5-PFV strand transfer that includes C3/Cy3-PFV bleed-through events (6%). The Gaussian fits to the histograms are shown as red lines. The TCC and STC data were collected at 100 msec and 1 sec resolutions, respectively.

The relative vDNA geometry in the Cy3/Cy5-PFV was examined during interactions with the model 1 nt Gap (5’-OH) target DNA that displayed the greatest TCC and STC formation. We first localized the 1 nt Gap (5’-OH) target DNA in the smFRET flow cell and then photobleached the identifying Cy5 fluorophore prior to injection of Cy3/Cy5-PFV intasomes. Transient TCC events similar to those observed with the Cy3-PFV intasomes were frequent and displayed a broad FRET peak with trailing less frequent lower FRET events (100 msec frame rate; *E*_*Cy3/Cy5-PFV*_ = 0.62 ± 0.1, mean ± 1*σ*; **Fig. 4c**; **Extended Data Fig. 12**). These observations suggest that formation of a TCC on 1 nt Gap (5’-OH) target DNA generally broadened the distance between the vDNAs compared to the distance calculated from the structural studies. Moreover, the broad FRET distribution is consistent with increased dynamic mobility of the intasome complex during target DNA binding.

In contrast, the formation of a stable STC by Cy3/Cy5-PFV intasomes on the 1 nt Gap (5’-OH) target DNA resulted in a narrower distribution of higher FRET events with fewer trailing lower FRET events (1 sec frame rate; *E*_*Cy3/Cy5-PFV*_ = 0.69 ± 0.06, mean ± 1*σ*; **Fig. 4d**). A similar narrower distribution with significantly fewer STC events was detected at 100 msec frame rate (*E*_*Cy3/Cy5-PFV*_ = 0.66 ± 0.09, mean ± 1*σ*; **Extended Data Fig. 12**). Both the TCC and STC events displayed a lower FRET efficiency than calculated from the crystal structures. This observation suggests that these intasome complexes are likely to exhibit a more open configuration in solution than proposed in the crystal structure ^22,30^. We also noted a peak of very low pseudo-FRET events that appeared when the studies were performed at 1 sec frame rate (6% of the products). When examined individually these pseudo-FRET events were found to result from STC events by Cy3/Cy3-PFV intasomes containing two Cy3-vDNAs. Excitation of these Cy3/Cy3-PFV STC events resulted in bleed-through to the Cy5 channel resulting in pseudo-FRET. When we added these to the total number of STC events (see: *) as well as the calculated frequency of Cy5/Cy5-PFV intasome STC events that are undetectable with this smFRET system (∼5%), the total frequency of STC events with these Cy3+Cy5 vDNA PFV intasomes (∼21%) is near equivalent to the frequency of STC events observed with the Cy3-PFV intasomes (20%). Overall, these results suggest that the STC events result in a relatively stable vDNA geometry on the target DNA. Since four-way DNA is dynamic even in the absence of nicks or gaps at the junction that should further increase mobility ^46^, these observations suggest that following STC formation a stable integrase complex remains associated with the covalent vDNA-target DNA structure.

## DISCUSSION

Previous single molecule analysis with PFV intasome demonstrated that recognition of a suitable DNA site limits integration ^23^. Once a DNA site is identified the concerted strand transfer steps occur in < 0.5 sec ^23^. Historical biochemical and structural studies have suggested that retroviruses prefer bent and/or flexible DNA at the integration site ^20,22,24,44^. Here we show that intrinsically flexible but intact duplex DNA containing a G/T mismatch or a +T nucleotide insertion that induces a stable DNA bend does not significantly enhance PFV intasome binding or integration. Interestingly, the substrates and structures that suggested a preference for bent and/or flexible DNA also altered the twist of the duplex target. We consider the possibility that it is the recognition of a stable or transiently untwisted duplex DNA segment that is the likely preferred retroviral intasome integration site for two reasons. First, structural studies of the PFV intasome STC with naked and nucleosome target DNA suggest strand transfer occurs across an untwisted backbone that averages approximately ∼12 bp per turn ^22,24^. Second, PFV clearly displays a preference for supercoiled DNA ^23^, which undergoes continuous and dynamic exchange between plectonemic supercoils and untwisted duplex DNA ^47^.

Nonetheless, we show that PFV intasomes also display a preference for target DNAs containing strand breaks and gaps, suggesting that bent DNA with many degrees-of-freedom not generally observed with intact duplex DNA can act as a binding sink. Moreover, this distinct binding enhances site-specific strand transfer, which appears to be the first targeted integration observed for any retrovirus. The smFRET analysis additionally detected an asymmetric TCC interaction with the nicked and gapped target DNA that places the PFV intasome on the 3’-side of the lesion. Integration site analysis correspondingly indicated asymmetric strand transfer to the 3’-side of the lesion. These results indicate that the PFV intasome stalls on the DNA next to the missing nucleotide (or break), where strand transfer into the lesion-containing strand is impossible because the phosphate bond is missing. Interestingly, this observation offers a plausible explanation for the 5’-G preference found in HIV integration site analysis, assuming the HIV intasomes are similarly targeted to nicked or gapped target DNA sites. In this model, retroviral intasomes pause asymmetrically at sites where OGG1 has created a gap following removal of an 8-oxo-G lesion along with associated 5’-dRP lyase activity. However, in the absence of a phosphate bond these targeted events are likely to result in half-site integration products that should result in a double strand break (DSB). To obtain a productive concerted integration, the phosphate bond must remain intact, or the gap must be repaired and followed by the second strand transfer before the intasome is disassembled or a DSB is formed. Interestingly, MUTYH has been found to interact with abasic sites shielding them from APE, thus preserving the phosphate bond ^48^. MUTYH also interacts with the RAD9-RAD1-HUS1 (9-1-1) DSB signaling and repair complex ^49^. Perhaps a combination of OGG1 and MUTYH activities creates a DNA lesion that targets retrovirus intasome binding as well as the components necessary to recover some concerted integration products from the largely half-site integration events. This notion would be consistent with the genetic studies that identified both *OGG1* and *MUTYH* as genes that affect retroviral integration ^8,9^. In this scenario, the loss of 5’-G preference in consensus HIV integration sites identified from *OGG1* mutant cells would be due to the loss of integration products that are targeted to 8-oxo-G lesions.

The smFRET conformational dynamics of Cy3/Cy5-PFV intasomes strongly suggest that the complex remains associated with the target DNA following an integration event (**Fig. 4**). Single molecule magnetic tweezer studies initially showed that under very low force (∼0.1 pN), any bound integration complex allows free rotation of DNA strands that released supercoils as well as dissociation of a DSB induced by a two-step strand transfer integration event with model vDNA oligonucleotides ^23^. Subsequent studies suggested that the intasome complex may be stable enough to tether the DSB ends even at high force (>30 pN) ^29^. However, these latter studies appear to leave the reaction on ice for extended periods of time prior to force induction. Under these conditions PFV intasomes undergo partial or complete activity loss as well as significant aggregation ^50^ that we observe here as background pseudo-FRET excursion events. It seems plausible that high force stability may largely reflect half-site integration products by partially inactivated intasomes that would be tethered by the unintegrated single stranded DNA and ultimately susceptible to mechanical breakage under these conditions. Nevertheless, our smFRET structural studies suggest that the IN subunits stabilize the vDNA geometry following integration albeit in a slightly more open configuration than predicted by the crystal structure.

Adaptation of genomic integration properties has been employed in the development of gene delivery vehicles designed to correct a variety of monogenic diseases ^51^. Lentiviral vectors derived from the HIV-1 retrovirus are most easily manipulated and can infect non-dividing cells ^52^. However, HIV-1 often integrates into open actively transcribing host genomic regions ^53^, which may unintentionally alter cellular functions or activate oncogenes ^54^. Modulating non-pathogenic retrovirus target site choice is one scheme for reducing the potential pathogenicity of genome targeting. The results presented here suggest for the first time that modifying the DNA configuration of a target site may be an additional method for targeted gene therapy.

## METHODS

Methods are included in a separate document for online publication.

## Acknowledgments

The authors would like to thank Ryan Messer and Yow-Yong Tan for technical assistance; and Rob Levendosky, and Gregory Bowman (John Hopkins University) for providing protocols for the sequencing ladder. This work was supported by the National Institutes of Health (AI126742 to R.F. and K.E.Y and GM121284 to K.E.Y).

## Author contributions

G.S., R.F., and K.E.Y. conceived, designed, and analyzed the smFRET experiments. A.G. performed and analyzed the integration site mapping experiments. J.L. wrote the MATLAB code for smFRET data analysis. G.S., A.G., J.L., R.F., and K.E.Y. wrote the manuscript.

## Additional information

Any supplementary information and source data are available in the online version of the paper. Correspondence and requests for materials should be addressed to K.E.Y. or R.F.

