## Supplementary material for "DNA Strand Breaks and Gaps Target Retroviral Binding and Integration": Methods

### ONLINE METHODS

#### Preparation of tDNAs

ssDNA DNA oligonucleotides used in this study are listed in **Extended Data Table 1**. The 8-Oxo G ssDNA was purchased from Midland Certified Reagent Company, Inc. All the other ssDNAs were purchased from Integrated DNA Technologies. ssDNAs were labeled with NSH-ester of A488 (GE Healthcare) or sulfo-Cy5 (Lumiprobe) at C6 amino modifications following the standard labeling protocol[1]. The labeled and unlabeled DNAs were separated on a C18 column (Agilent Technologies) using reverse-phase HPLC. Selected HPLC fractions were concentrated using 0.5 mL, 3 kDa molecular weight cutoff Amicon Ultra centrifugal filters after evaporating the organic solvent in a SppedVac vacuum concentrator (Thermo Fisher Scientific). The DNAs were buffer exchanged into 20 mM Tris-HCl pH 8.0, 1 mM EDTA, and stored in -20 $^0$C until further purification.

Labeled ssDNAs prepared in this manner still significantly contaminated with truncations from the synthesis. Therefore, each ssDNA was gel purified to ~100% purity using 12% Acrylamide:Bis 19:1/7M Urea PAGE. The gel extracted DNAs were concentrated as above and stored at -20 $^0$C in 20 mM Tris-HCl pH 8.0, 1 mM EDTA.

The substrate DNAs in **Extended Data Table 1** were obtained by annealing equimolar mixtures of corresponding ssDNAs in 20 mM Tris-HCl pH 8.0, 1 mM EDTA, 100 mM NaCl. Annealing reactions were done in a thermal cycler (Thermo Fisher Scientific) by heating the samples to 95 $^0$C and slowly cooling down to 15 $^0$C. Fully annealed dsDNAs were enriched by ion-exchanging HPLC on a Gen-Pak Fax column (Waters). The purity of HPLC fractions was tested by 5% Acrylamide:Bis 59:1 native PAGE. Fractions containing ~100% pure dsDNA were pooled together and concentrated as above. The concentrations of dsDNAs were determined by UV-Vis absorbance of the DNA at 280 nm, AF488 at 490 nm, and Cy5 at 650 nm using a nanodrop device (Thermo Fisher Scientific). These DNAs were stored at -20 $^0$C in 20 mM Tris-HCl pH 8.0, 1 mM EDTA, 100 mM NaCl.

**PFV intasome assembly**

PFV intasomes were assembled as described previously using recombinant integrase and dsDNA mimicking the U5 vDNA ends (**Extended Data Table 1**)[2-4]. vDNAs were prepared as described above using appropriate ssDNA oligos (**Extended Data Table 1**). The intasome assemblies were performed by salt dialysis followed by chromatographic purification on a size exclusion column (SEC). The activities of SEC fractions were tested using a standard plasmid-based integration assays before flash freezing and storing at -80 $^0$C for further use. Catalytically deficient Cy3-PFV-ddA construct was assembled in the same way using integrase and a vDNA containing 3'-ddA (**Extended Data Table 1**)[5,6]. The intasome containing the Cy3 and Cy5 FRET pair (Cy3/Cy5-PFV) was prepared by mixing an equimolar Cy3 and Cy5 vDNAs during the assembly reaction. The expected outcomes for different species are: 50% Cy3 and Cy5 containing intasomes, 25% two Cy3 intasomes, and 25% two Cy5 intasomes.

**$R_{Cy3\text{-}Cy5}$ and $E_{TCC\ or\ STC}$ estimations from the crystal structures**

The PFV intasome TCC and STC crystal structures (PDB code: 3OS1, 3OS0) were downloaded from the PDB data base. To measure the distances ($R_{Cy3\text{-}Cy5}$) between the fluorophores in vDNAs and the target DNAs, we extended the target DNA chains in both crystal structures due to absence of some base pairs in the structures compared to the target DNA used in the smFRET experiments. This was done by loading the structures into the discover studio and generating a B-helix of double strands DNA structures (sequence: CCCGAG), which then ligated to the ends of the target DNA. The resultant structures were then opened in PyMol 2.1 (Schrödinger, Inc) to estimate inter dye distances ($R_{Cy3\text{-}Cy5}$) for various complexes. Although the dyes were attached through C6 amino linkers, the $R_{Cy3\text{-}Cy5}$ were measured between DNA bases using the distance measurement module in PyMol. These $R_{Cy3\text{-}Cy5}$ were converted to corresponding FRET efficiencies using the Förster equation, assuming the orientation factor ($\kappa^2$) is 2/3 (complete free rotations of the dyes) and the Förster radius for Cy3-Cy5 ($R_{0,Cy3\text{-}Cy5}$) is 6 nm[7].

$$E = \frac{1}{1+\left(\frac{R_{Cy3-Cy5}}{R_{0,Cy3-Cy5}}\right)^6} \quad (1)$$

For the construct that showed two different FRET state because of the presence of two FRET pairs the expected FRET ($E_{TCC}$ or $E_{STC}$) was determined as arithmetic mean of the two states.

**smFRET imaging**

smFRET imaging was done on a home built inverted fluorescence microscope (Olympus), as described previously[1]. Prism-based total internal reflection (TIRF) of a green (532 nm) or red (635 nm) laser was used to excite fluorophores attached to a surface of flow cell. The fluorescence from individual fluorophores was collected through a 60X water immersion objective (Olympus) and directed onto an emCCD chip (Princeton Instruments) after magnifying another 1.6X and separating Cy3, Cy5 emissions using a Dual View device (Photometrics).

The quartz surface of the flow cells was passivated with a 1:20 ratio of biotin-PEG and methoxy-PEG (5000 MW, Layson Bio, Inc). Biotin-PEG was used to immobilize target DNAs by biotin-neutravidin-biotin interactions at ~0.2 molecules/μm$^2$ surface density. The methoxy-PEG brush minimizes the surface interactions of biomolecules[1,7]. The imaging buffer (Buffer-I) for all the experiments consisted of 30 mM Bis-tris propane, pH 7.5, 110 mM NaCl, 2 mM $MgSO_4$, 4 μM $ZnCl_2$, 0.1 mM DTT, 0.2 mg/mL BSA, 0.02% IGPEPAL. Buffer-I also included a cocktail of saturated (~2 mM) Trolox and an oxygen scavenging system (OSS) to minimize photo-blinking and photobleaching of fluorophores respectively[1]. The OSS consisted of 25 mM protocatechuic acid (PCA) and 20 nM protocatechuate dioxygenase (PCD)[1,8]. All the experiments were performed at 24 ± 1 $^0$C.

**smFRET target capture assays**

The imaging for target capture was done at 100 ms time resolution to capture transient events. Single-molecule movies were initiated by exciting Cy5-DNA in Buffer-I with a 635 nm red laser at ~2 mW. After 20 s, the excitation was switched to a 532 nm green laser maintained at ~6 mW. 10 s after the green laser exposure, 5 nM Cy3-PFV in Buffer-I was infused in real-time into the flow cells. Data recording was continued under continuous green laser excitation for 2.5 min from the injection (**Extended Data Fig. 2a**).

Cy3/Cy5-PFV experiments were performed the same way with the following modifications. The initial red laser exposure was used to bleach Cy5 in the field of view (FOV) entirely within 20 s. The fast photobleaching was achieved by eliminating the OSS in Buffer-I. The subsequent intasome injection and imaging were done exactly as above with OSS in Buffer-I (**Extended Data Fig. 2b**).

**smFRET strand transfer assays**

Strand transfer assays were recorded with 1 s time resolution and at lower laser powers to improve fluorophore lifetimes, allowing longer observations. Movies were initiated by Cy5 excitation with the red laser at ~1 mW. After 30 s, the excitation was switched to the green laser maintained at ~4 mW. 30 s after the green laser exposure, 5 nM Cy3-PFV in Buffer-I was infused in real-time into the flow cell. The imaging was continued under continuous green laser excitation for 10 min from the injection (**Extended Data Fig. 2c**).

Cy3/Cy5-PFV experiments were performed the same way with the following modifications. The initial red laser exposure was used to bleach Cy5 in the FOV entirely within 30 s. The fast photobleaching was achieved by eliminating the OSS in Buffer-I. The subsequent intasome injection and imaging were done exactly as above with OSS in Buffer-I (**Extended Data Fig. 2d**).

The Rev-Cy5 1nt Gap (5'-OH) experiments suffer from fast photobleaching and photophysical fluctuations of Cy5 due to its closeness to purines (A,G)[9] in the DNA sequence

(**Extended Data Table 1**). Therefore, we performed only 1 s resolution experiments at ~2 mW reduced power for 532 nm laser in regular experiments and at ~4 mW for fast photobleaching.

**Initial processing of SM movies**

Single-molecule movies were acquired using the Micro-Manager imaging software as described previously (**Supplementary Movies 1-7**)[1]. We used a custom-written MATLAB (MathWorks) program to extract intensity and FRET data from these movies as follows.

Prior to the analysis the program requires mapping of the Cy3 and Cy5 channels. This was accomplished by using the emissions of 0.2 μm crimson carboxylate modified microspheres (Thermo Fisher Scientific). Then using the program, the movies were manually inspected to identify single-molecules as diffraction-limited spots. The initial Cy5 excitation provided excellent signal to noise (S/N) for identifying individual DNA molecules. Each molecule is then marked with a circle of adjustable radius. However, the crowding of DNAs in the FOV may produce overlapping circles. To avoid interference, we chose smaller three-pixel radius that encircled an entire molecule without including neighbors. In addition, we set a five pixels distance cutoff from a molecule to another or to the edges of the FOV. The eccentricity (a measure of circularity) cutoff for the spots was chosen as 0.2. All these stringent selection criteria allowed us to identify ~500-600 well separated DNAs per movie.

**Generation of SM traces and HMM analysis**

After the selection algorithm, pixel intensities within a given circle were integrated to generate raw Cy3 and Cy5 intensities. This procedure was continued for all the movie frames to collect emissions as a function of time and to build intensity traces without background corrections **(Extended Data Fig. 2a-d, panel 1).** The graphical user interface (GUI) of our MATLAB program allowed direct comparison of these traces with their corresponding overlapped Cy3, Cy5 spots in the single-molecule movies.

The infusion of Cy3-PFV led to marked increase in both Cy3 and Cy5 backgrounds. These intensity jumps were used as references for trace truncations and background corrections **(Extended Data Fig. 2a-d, panel 2).** The 100 ms traces were smoothed with a three-point averaging and the 1 s traces were not smoothed. Truncated traces were subjected to vbFRET HMM algorithm built into our program[10] to first adjust the backgrounds and then to identify states in FRET traces. FRET was calculated from corrected intensities (I) as $I_{Cy5}/(I_{Cy3}+I_{Cy5})$. Four states were used as the initial guess for HMM. When an intasome is not bound to a target DNA, both $I_{Cy3}$ and $I_{Cy5}$ approached zero in the background corrected intensity traces. This led to erratic fluctuations in FRET for intasome unbound regions **(Extended Data Fig. 2a-d, panel 3).** As a solution, the FRET and the corresponding HMM state was assigned zero when either $I_{Cy3}$ or $I_{Cy5}$ approached zero. These 'cleaned up' FRET traces and their HMM fittings were manually inspected for correct fittings **(Extended Data Fig. 2a-d, panel 4).** Moreover, traces containing Cy3 photobleaching indicated negligible spectral bleed-through to the Cy5 channel. Therefore, Cy3 bleed-though correction was omitted from $I_{Cy5}$. Also, intasome aggregates with saturating Cy3 only contributed to a minor Cy5 signal and 0.06 FRET.

**Further selection and categorization of smFRET traces**

Intasome injections sometimes created photophysical fluctuations at the beginning of some of the FRET traces. When necessary these aberrant data were excluded by truncating the traces (up to 100 frames). In other cases, the surface binding of an intasome within a target DNA selection circle resulted in steady Cy3 signals without DNA binding. However, visual inspection of movies allowed us to identify these pseudo-events and elimination by truncation. Only molecules that contained at least 100 frames worth data were included in the final analyses.

Traces containing FRET or colocalized Cy3-Cy5 for prolonged time windows (up to minutes) were categorized as STCs. Traces that lack these long events were categorized as TCCs. The data for these two categories were analyzed and presented separately. $\%STC_{FRET}$

and their error estimates ($\Delta\%STC_{FRET}$) were calculated from the number of DNAs that showed strand transfer events (*n*) and total number of DNA molecules (*N*), using equation (2) and (3) respectively[11].

$$\%STC_{FRET} = {}^{n}/_{N} \times 100\% \tag{2}$$

$$\Delta\%STC_{FRET} = {}^{\sqrt{n}}/_{N} \times 100\% \tag{3}$$

**Transition Density Plots (TDPs) for target capture smFRET traces**

Using our MATLAB program TDPs were generated by compiling idealized FRET traces resulting from the HMM fittings[10]. The peaks in a TDP represent transitions from a given initial FRET state to a given final FRET state. Also, the peak heights represent total number of transitions between corresponding states[10]. The use of 4 states as an initial guess for HMM, resulted in occasional overfitting of the data and low populated off diagonal peaks.

**Post-Sync Histogram analysis**

Using our MATLAB program FRET traces were synchronized to the injection of intasomes to create PSH plots. The assigned 0 FRET corresponding to the intasome unbound state was eliminated for clarity. For 100 ms resolution experiments, 0.01 FRET bins and 300 ms time bins were used to construct PSHs. For 1 s experiments, 0.01 FRET bins and 0.91 s time bins were used to construct PSHs.

**smFRET histograms**

smFRET histograms were built with 0.2 FRET bins by molecule and time-averaging the >0 FRET states in the traces. The heights of the bins (counts) depend on the number of target DNA molecules included in the analysis. For TCC histograms, this dependency was eliminated by recalculating the counts per DNA. To directly compare between Cy3-PFV experiments the histograms were re-normalized to the number of events in the pseudo-FRET ~0.06 peak. This was done by integrating the area under the pseudo-FRET peak to calculate the raw counts and then dividing the whole histogram by that number.

All the smFRET histograms were prepared and fitted using Igor Pro 8 (WaveMetrics). The following Gaussian equation was used for fittings.

$$y = \sum_{i=1}^{n} 1 / \sqrt{2\pi\sigma_i^2}\, e^{\frac{(x-x_{0,i})^2}{2\sigma_i^2}} \qquad (4)$$

In most cases a single Gaussian (n =1) fit the data well. Occasionally n = 2 or n = 3 were needed to get the best fits.

**Dwell time and transition count histograms**

The dwell times of the bound ($t_{on}$), and unbound ($t_{off}$) states were extracted from the TCC traces using the HMM fittings, as shown in **Extended Data Fig. 8a**. For Cy3-PFV intasome experiments, using a FRET threshold of 0.1, the FRET > 0.1 was defined as bound, and 0 FRET was defined as unbound (**Extended Data Fig. 8a**). The dwell times from many molecules were binned according to the web-based bin optimization algorithm (https://www.neuralengine.org//res/histogram.html), and histograms were generated in Igor Pro 8 (WaveMetrics). Following single exponential functions were used to calculate the average bound ($\tau_{on}$) and unbound ($\tau_{off}$) dwell times.

$$Counts_{bound} = A_1 e^{-t_{on}/\tau_{on}} \qquad (5)$$

$$Counts_{unbound} = A_2 e^{-t_{off}/\tau_{off}} \qquad (6)$$

The transition counts for Cy3-PFV TCC traces were calculated using FRET thresholds; 0.1. Upward FRET jumps that crossed the threshold was defined as transitions. Transition counts from individual traces were binned with 1-transition bins to build histograms in Igor pro 8 (WaveMetrics).

**Overall ST times**

Strand transfer times ($t_{ST}$) is defined as the time from the intasome injection to the first frame of that a stable FRET or Cy3 signal appear (**Extended Data Table 6**). A MATLAB script was used to extract $t_{ST}$ from individual traces. The mean ($\bar{\boldsymbol{t}}_{\boldsymbol{ST}}$) and standard deviation ($\boldsymbol{\sigma}_{\boldsymbol{ST}}$ .) for a given target DNA was calculated as,

$$\bar{t}_{ST} = \frac{\sum_{i=1}^{N} t_{ST}}{n} \tag{7}$$

$$\boldsymbol{\sigma}_{\boldsymbol{ST}} = \sqrt{\frac{1}{n-1}\sum_{i=1}^{n}(t_{ST} - \bar{t}_{ST})^2} \tag{8}$$

Where *n* is the number DNA molecules that showed strand transfer events.

**Ensemble biochemical experiments**

**Plasmid-based integration assay**

This assay was performed using the standard protocol described previously[2,3]. Briefly, 25 nM of PFV INTs were incubated with 50 ng of supercoiled (SC) plasmid DNA (pGEM®-T Easy) in 10 mM Bis-tris propane, pH7.5, 110 mM NaCl, 5 mM $MgSO_4$, 4 µM $ZnCl_2$ 10 mM DTT in 15 µl reactions for 5 min at 37 $^0$C. The reactions were terminated by adding 0.1% SDS, 2.5 mM EDTA, 1 mg/ml proteinase K and incubated at 55 $^0$C for an hour. The products were mixed with 5% glycerol before resolving on a 1% agarose gel in 1X TAE at 105 V for an hour. Gels were stained with 0.1 µg/mL ethidium bromide and scanned on a Sapphire Biomolecular Imager. The quantification of supercoiled (SC) and linear DNA was done using the AzureSpot software (Azure Biosystems) as described previously[2,3] and presented in **Extended Data Fig. 1**.

**Integration site mapping experiments**

Integration site mapping experiments were also performed as described previously[4] using target DNAs in **Extended Data Table 2**. Briefly, 10 nM unlabeled PFV intasomes were incubated with

5 nM targets in 30 mM Bis-tris propane, pH7.5, 110 mM NaCl, 2 mM $MgSO_4$, 4 μM $ZnCl_2$, 10 mM DTT in 15 μl reactions for 5 min at 37 °C. Reactions were terminated by adding 0.1% SDS, 2.5 mM EDTA, 1 mg/ml proteinase K and incubating at 37°C for 20 min. Deproteinated samples were denatured by heating to 95 °C with 50% formamide for 10 min, then iced. Sequencing ladders were generated using Thermo Sequenase Dye Primer Manual Cycle Sequencing Kit 792601 according to the manufacturer's directions[12] with pDrive-601NPS[1] as the template and Cy5, Alexa488 or Cy3 labeled oligos complementary to the 5' end of 601NPS as primers. 1 pmol primer and I pmol template per reaction were used. Annealing temperature was 55 °C with 45 cycles. Sequencing reactions were diluted with an equal volume of formamide, heated to 75 °C for 10 min and stored at -20 °C. Products were resolved on 0.8 mm 8% Acrylamide:Bis 19:1/7M Urea PAGE gels in 1XTBE at 40 W for various times. Gels were scanned on a Sapphire Biomolecular Imager and quantified using AzureSpot software (Azure Biosystems). Alignment of the products with the sequencing ladder was used to determine the integration sites. The integration efficiency was calculated as the fractional intensity of a band relative to the lane. Reactions without INTs were used as controls for target input.

The site mapping experiments comparing different 1nt Gap (5'-OH) DNAs were performed as described above with the following modifications in the reaction conditions. An intasome containing shorter vDNAs and Cy3 at the transfer strand (Cy3-PFV(TS), **Extended Data Table 1**) at 100 nM was reacted with 25 nM DNA targets to improve the efficiency of the reaction. In addition, the reaction buffer was supplemented with 5 nM PCA to improve the lifetime of the intasome[3]. The PAGE separation of resultant products, the site size determination and imaging were done same as above.
