## Extended Data for "DNA Strand Breaks and Gaps Target Retroviral Binding and Integration"

**Extended Data Table 1. Target DNAs and vDNAs Used in this Study.**

| Oligo names | Sequence | DNA Substrate |
|---|---|---|
| KEY22<br>KEY21 | 5′-Bio-CTGGAGAATCCCGGTGCCGAGGCCGCTCAATTGGTCGTAGACAGCTCTAGCACCGCTTAA-3′<br>3′-GACCTCTTAGGGCCACGGCTCCGGCGAGTTAACCAGCATCTGTCGAGATCGTGGCGAATT-5′ | GC |
| KEY25<br>KEY21 | 8-OxoG<br>5′-Bio-CTGGAGAATCCCGGTGCCGAGGCCGCTCAATTGGTCGTAGACAGCTCTAGCACCGCTTAA-3′<br>3′-GACCTCTTAGGGCCACGGCTCCGGCGAGTTAACCAGCATCTGTCGAGATCGTGGCGAATT-5′ | 8-OxoG |
| KEY22<br>KEY28 | 5′-Bio-CTGGAGAATCCCGGTGCCGAGGCCGCTCAATTGGTCGTAGACAGCTCTAGCACCGCTTAA-3′<br>3′-GACCTCTTAGGGCCACGGCTCCGGCGAGTTAACCAGCATTTGTCGAGATCGTGGCGAATT-5′ | G/T |
| KEY29<br>KEY21 | T<br>5′-Bio-CTGGAGAATCCCGGTGCCGAGGCCGCTCAATTGGTCGTAGACAGCTCTAGCACCGCTTAA-3′<br>3′-GACCTCTTAGGGCCACGGCTCCGGCGAGTTAACCAGCATCTGTCGAGATCGTGGCGAATT-5′ | +T |
| KEY23 KEY24 (5'-P)<br>KEY 21 | HO P<br>5′-Bio-CTGGAGAATCCCGGTGCCGAGGCCGCTCAATTGGTCGTAGACAGCTCTAGCACCGCTTAA-3′<br>3′-GACCTCTTAGGGCCACGGCTCCGGCGAGTTAACCAGCATCTGTCGAGATCGTGGCGAATT-5′ | Nick (5'-P) |
| KEY23 KEY26 (5'-P)<br>KEY 21 | HO P<br>5′-Bio-CTGGAGAATCCCGGTGCCGAGGCCGCTCAATTGGTCGTA ACAGCTCTAGCACCGCTTAA-3′<br>3′-GACCTCTTAGGGCCACGGCTCCGGCGAGTTAACCAGCATCTGTCGAGATCGTGGCGAATT-5′ | 1nt Gap (5'-P) |
| KEY27 KEY26 (5'-P)<br>KEY 21 | HO P<br>5′-Bio-CTGGAGAATCCCGGTGCCGAGGCCGCTCAATTGGTCGT ACAGCTCTAGCACCGCTTAA-3′<br>3′-GACCTCTTAGGGCCACGGCTCCGGCGAGTTAACCAGCATCTGTCGAGATCGTGGCGAATT-5′ | 2nt Gap (5'-P) |
| KEY23 KEY24 (5'-OH)<br>KEY 21 | HO OH<br>5′-Bio-CTGGAGAATCCCGGTGCCGAGGCCGCTCAATTGGTCGTAGACAGCTCTAGCACCGCTTAA-3′<br>3′-GACCTCTTAGGGCCACGGCTCCGGCGAGTTAACCAGCATCTGTCGAGATCGTGGCGAATT-5′ | Nick (5'-OH) |
| KEY23 KEY26 (5'-OH)<br>KEY 21 | HO OH<br>5′-Bio-CTGGAGAATCCCGGTGCCGAGGCCGCTCAATTGGTCGTA ACAGCTCTAGCACCGCTTAA-3′<br>3′-GACCTCTTAGGGCCACGGCTCCGGCGAGTTAACCAGCATCTGTCGAGATCGTGGCGAATT-5′ | 1nt Gap (5'-OH) |
| KEY27 KEY26 (5'-OH)<br>KEY 21 | HO OH<br>5′-Bio-CTGGAGAATCCCGGTGCCGAGGCCGCTCAATTGGTCGT ACAGCTCTAGCACCGCTTAA-3′<br>3′-GACCTCTTAGGGCCACGGCTCCGGCGAGTTAACCAGCATCTGTCGAGATCGTGGCGAATT-5′ | 2nt Gap (5'-OH) |
| KEY23 Cy5-KEY26 (5'-OH)<br>KEY 21 | HO OH<br>5′-Bio-CTGGAGAATCCCGGTGCCGAGGCCGCTCAATTGGTCGTA ACAGCTCTAGCACCGCTTAA-3′<br>3′-GACCTCTTAGGGCCACGGCTCCGGCGAGTTAACCAGCATCTGTCGAGATCGTGGCGAATT-5′ | R-Cy5<br>1nt Gap (5'-OH) |
| KEY675<br>Cy3-KEY616 | 5′-CTGTTCGGGCGCCACTCAATATACAAAATTCCATGACA-3′<br>3′-GACAAGCCCGCGGTGAGTTATATGTTTTAAGGTACTGTTA-5′ | Cy3-PFV |
| 3'-ddA KEY675<br>Cy3-KEY616 | 5′-CTGTTCGGGCGCCACTCAATATACAAAATTCCATGAC[2′3′ddA]-3′<br>3′-GACAAGCCCGCGGTGAGTTATATGTTTTAAGGTACTGTTA-5′ | Cy3-PFV-ddA |
| KEY675<br>KEY616 | 5′-CTGTTCGGGCGCCACTCAATATACAAAATTCCATGACA-3′<br>3′-GACAAGCCCGCGGTGAGTTATATGTTTTAAGGTACTGTTA-5′ | PFV |
| Cy3-KEY675(2)<br>KEY616 | 5′-GCGCCACTCAATATACAAAATTCCATGACA-3′<br>3′-CGCGGTGAGTTATATGTTTTAAGGTACTGTTA-5′ | Cy3-PFV (TS) |
| KEY675<br>Cy5-KEY616 | 5′-CTGTTCGGGCGCCACTCAATATACAAAATTCCATGACA-3′<br>3′-GACAAGCCCGCGGTGAGTTATATGTTTTAAGGTACTGTTA-5′ | Cy5-PFV |

The constituent DNA oligonucleotides, the target DNA substrates, and the vDNAs used in this study. The labeling positions of A488, Cy3 and Cy5 are marked with blue, green, and red colors respectively. Bio = 5'-Biotin, HO = Hydroxyl, P = Phosphate.

**Extended Data Table 2. $E_{TCC}$ and $E_{STC}$.**

| DNA Substrate | $E_{TCC} \pm \sigma_{TCC}$ | $E_{STC} \pm \sigma_{STC}$ |
|---|---|---|
| GC | – | 0.54 ± 0.06 |
| G/T | – | 0.32 ± 0.08 |
| Nick (5'-P) | 0.15 ± 0.08 | 0.27 ± 0.04, 0.10± 0.04 |
| 1nt Gap (5'-P) | 0.16 ± 0.05 | 0.16 ± 0.05 |
| 2nt Gap (5'-P) | 0.18 ± 0.10 | 0.21 ± 0.05 |
| Nick (5'-OH) | 0.17 ± 0.07 | 0.17 ± 0.05 |
| 1nt Gap (5'-OH) | 0.19 ± 0.05 | 0.18 ± 0.05 |
| 2nt Gap (5'-OH) | 0.23 ± 0.11 | 0.24 ± 0.05 |
| Blocked end 1nt Gap (5'-OH) | 0.18 ± 0.05 | 0.19 ± 0.05 |
| 1nt Gap (5'-OH) + Cy3-PFV-ddA | 0.15 ± 0.05 | - |
| R-Cy5 1nt Gap (5'-OH) | 0.21 ± 0.10 | 0.29 ± 0.06 |

These values were calculated as described in **ONLINE METHODS** by fitting TCC and STC FRET distributions for each experiment with a single or a combination of two Gaussians. *E* and $\sigma$ indicate the centers and the standard deviations of the peaks respectively.

**Extended Data Table 3. Statistics for Pseudo-FRET from the TCC Histograms Recorded at 100 ms Frame Rate..**

| DNA Substrate | Number of DNA molecules (N) | Pseudo FRET | Pseudo FRET |
|---|---|---|---|
| | | Total number of counts (n) | Number of counts per DNA (n/N) |
| GC | 552 | 283 | 0.513 |
| 8-OxoG | 496 | 258 | 0.521 |
| G/T | 574 | 342 | 0.596 |
| +T | 606 | 442 | 0.730 |
| Nick (5'-P) | 549 | 314 | 0.571 |
| 1nt Gap (5'-P) | 540 | 381 | 0.706 |
| 2nt Gap (5'-P) | 474 | 322 | 0.680 |
| Nick (5'-OH) | 558 | 346 | 0.621 |
| 1nt Gap (5'-OH) | 496 | 356 | 0.718 |
| 2nt Gap (5'-OH) | 540 | 194 | 0.359 |
| Blocked end 1nt Gap (5'-OH) | 483 | 297 | 0.614 |

$$\bar{\boldsymbol{x}} = 0.603, \sigma = \pm 0.110$$

These numbers were calculated as described in **ONLINE METHODS** by fitting FRET distributions for each DNA with a single or a combination of two Gaussians and integrating the area under the curve for the 0.06 FRET peak. $\bar{\boldsymbol{x}}$ and $\sigma$ show the average and the standard deviation for the counts per DNA calculated from all the DNA substrates.

**Extended Data Table 4. Statistics for Pseudo-FRET from the TCC Histograms Recorded at 1 s Frame Rate.**

| DNA Substrate | Number of DNA molecules (N) | Pseudo FRET | Pseudo FRET |
|---|---|---|---|
| | | Total number of counts (n) | Number of counts per DNA (n/N) |
| GC | 583 | 606 | 1.04 |
| 8-OxoG | 624 | 2359 | 3.78 |
| G/T | 613 | 1765 | 2.88 |
| +T | 531 | 1529 | 2.88 |
| Nick (5'-P) | 583 | 2268 | 3.89 |
| 1nt Gap (5'-P) | 542 | 2000 | 3.69 |
| 2nt Gap (5'-P) | 498 | 1604 | 3.22 |
| Nick (5'-OH) | 614 | 1105 | 1.8 |
| 1nt Gap (5'-OH) | 489 | 2308 | 4.72 |
| 2nt Gap (5'-OH) | 534 | 951 | 1.78 |

$\bar{x} = 2.97, \sigma = \pm 1.14$

These numbers were calculated as described in **ONLINE METHODS** by fitting FRET distributions for each DNA with a single or a combination of two Gaussians and integrating the area under the curve for the 0.06 FRET peak. $\bar{x}$ and $\sigma$ show the average and the standard deviation for the counts per DNA calculated from all the DNA substrates.

**Extended Data Table 5. Frequency (%)[1] of STC Events.**

| Cy3-PFV | | | PFV |
|---|---|---|---|
| **DNA Substrate** | **$\%STC_{smFRET}$ ± $\Delta\%STC_{smFRET}$** | **Number of DNA molecules (N)** | **$\%STC_{Gel}$ ± $\Delta\%STC_{Gel}$** |
| GC | 0.3 ± 0.2 | 585 | 0.4 ± 0.6 |
| 8-OxoG | 0 ± 0 | 624 | 0.9 ± 1.5 |
| G/T | 0.5 ± 0.3 | 616 | 1.4 ± 0.8 |
| +T | 0 ± 0 | 531 | 0.8 ± 0.7 |
| Nick (5'-P) | 0.7 ± 0.3 | 587 | 3.2 ± 2.9 |
| 1nt Gap (5'-P) | 3 ± 0.7 | 559 | 8.4 ± 3.8 |
| 2nt Gap (5'-P) | 1.4 ± 0.5 | 506 | 10.7 ± 5.7 |
| Nick (5'-OH) | 2 ± 0.6 | 626 | 20 ± 12.4 |
| 1nt Gap (5'-OH) | 20 ± 1.8 | 614 | 29.3 ± 10.9 |
| 2nt Gap (5'-OH) | 12 ± 1.4 | 605 | 14.2 ± 1.7 |
| Blocked end 1nt Gap (5'-OH) | 24 ± 2.0 | 547 | |
| R-Cy5 1nt Gap (5'-OH) | 20 ± 1.8 | 527 | |
| **DNA Substrate** | **$\%STC_{smFRET}$ ± $\Delta\%STC_{smFRET}$** | **Number of DNA molecules (N)** | |
| GC | 0 ± 0 | 547 | |
| 1nt Gap (5'-P) | 5 ± 0.9 | 579 | |
| 1nt Gap (5'-OH) | 22 ± 2.0 | 569 | |
| **Cy3-PFV-ddA** | | | |
| **DNA Substrate** | **$\%Long_{smFRET}$ ± $\Delta\%Long_{smFRET}$** | **Number of DNA molecules (N)** | |
| 1nt Gap (5'-OH) | 0.4 ± 0.2 | 564 | |
| **Cy3/Cy5-PFV** | | | |
| **DNA Substrate** | **$\%STC_{smFRET}$ ± $\Delta\%STC_{smFRET}$** | **Number of DNA molecules (N)** | |
| GC | 0.4 ± 0.3 | 548 | |
| 1nt Gap (5'-OH) | 16 ± 1.6 [2] | 609 | |

[1] Percentages were calculated as described in **ONLINE METHODS**. For Cy3-PFV-ddA this number was defined with a different name to signify the catalytic deficiency. *N* indicates the total number of DNA molecules analyzed for each experiment. *n* is the number of molecules that showed strand transfer events. The errors ($\Delta\%STC_{smFRET}$) reflect $\frac{\sqrt{n}}{N} \times 100\%$.

[2] Includes PFV intasomes containing Cy5-only integration products. Integration by PFV intasomes containing Cy3-only would not be detected in this smFRET system and could theoretically introduce an additional 5% STC, which would be virtually identical to 1 nt Gap (5'-OH) STC events (above).

**Extended Data Table 6. Strand Transfer Times.**

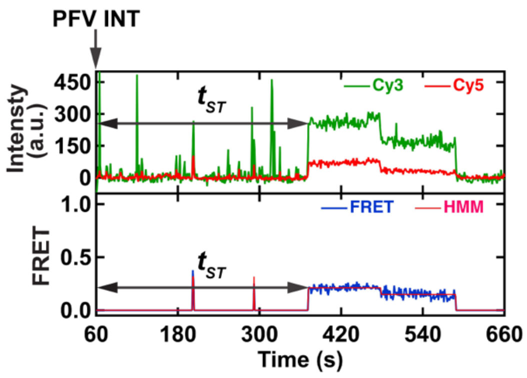

| **Cy3-PFV** | | |
|---|---|---|
| **DNA Substrate** | $\bar{t}_{ST} \pm \sigma_{ST}$ **(s)** | **Number of events (n)** |
| GC | 172 ± 87 | 2 |
| G/T | 328 ± 228 | 3 |
| Nick (5'-P) | 191 ± 143 | 4 |
| 1nt Gap (5'-P) | 254 ± 202 | 17 |
| 2nt Gap (5'-P) | 264 ± 176 | 7 |
| Nick (5'-OH) | 180 ± 147 | 12 |
| 1nt Gap (5'-OH) | 249 ± 150 | 125 |
| 2nt Gap (5'-OH) | 277 ± 148 | 71 |
| Blocked end 1nt Gap (5'-OH) | 260 ± 145 | 134 |
| R-Cy5 1nt Gap (5'-OH) | 216 ± 152 | 104 |
| **Cy3-PFV (TS1)** | | |
| **DNA Substrate** | $\bar{t}_{ST} \pm \sigma_{ST}$ **(s)** | **Number of events (n)** |
| 1nt Gap (5'-P) | 229 ± 166 | 28 |
| 1nt Gap (5'-OH) | 255 ± 159 | 124 |
| **Cy3-PFV-ddA** | | |
| **DNA Substrate** | $\bar{t}_{long} \pm \sigma_{long}$ **(s)** | **Number of events (n)** |
| 1nt Gap (5'-OH) | 186 ± 103 | 2 |
| **Cy3/Cy5-PFV** | | |
| **DNA Substrate** | $\bar{t}_{ST} \pm \sigma_{ST}$ **(s)** | **Number of events (n)** |
| GC | 140 ± 38 | 2 |
| 1nt Gap (5'-OH) | 251 ± 156 | 95 |

The strand transfer time ($t_{ST}$) is the time it takes to observe a stable FRET (or Cy3) signal from the intasome injection, as shown by the figure in the first raw. The mean values ($\bar{t}$) and standard deviations ($\sigma$) were calculated as described in **Online Methods**. For Cy3-PFV-ddA, this number was defined with a different name to signify the catalytic deficiency. *n* indicates the number of DNAs that showed strand transfer events.

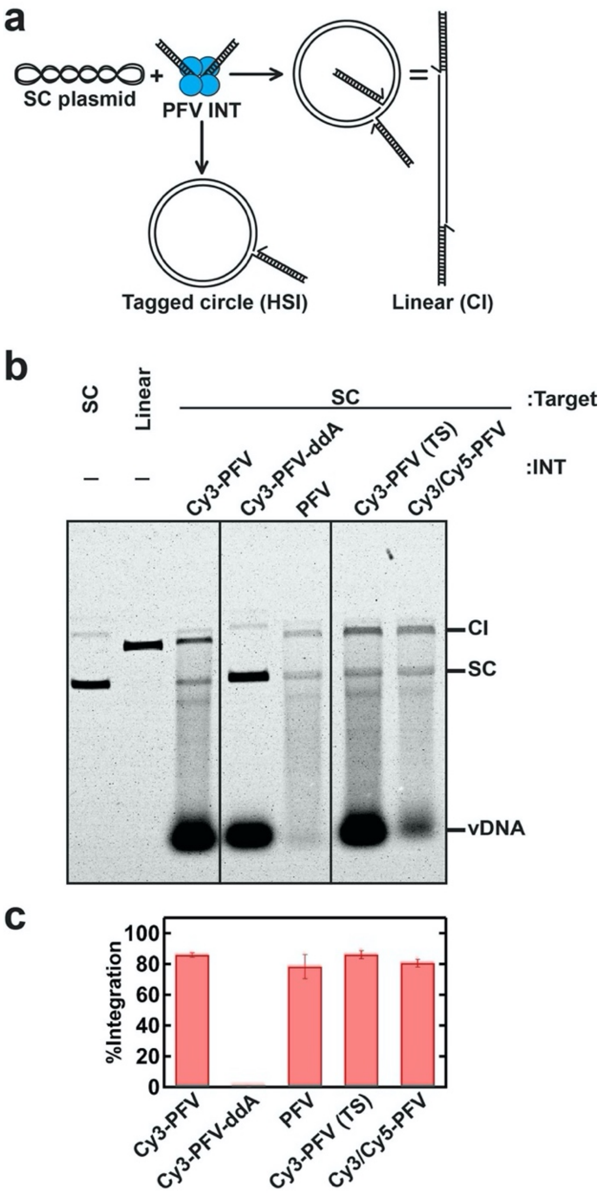

**Extended Data Figure 1. Integration Activity of the PFV Intasomes used in these Studies.** (**a**) An illustration of the gel-based analysis. When a PFV intasome consisting of DNA oligonucleotides mimicking vDNA ends catalyzes concerted integration into a supercoiled (SC) DNA, linear DNA is produced. A relaxed tagged circle is created when only one strand transfer occurs in a half-site integration (HIS). (**b**) An ethidium bromide-stained agarose gel showing the integration activity of all the intasomes used in this study. Competing aggregation and auto integration reactions continuously deplete the active pool of intasomes available for the reaction and cause the incomplete conversion of the SC DNA. (**c**) Quantification of the gel data using the ethidium bromide fluorescence. Error bars indicate standard deviations from two independent experiments. HSI products that should migrate slower than the linear DNA were undetectable with ethidium bromide. The catalytically deficient Cy3-PFV-ddA did not produce any detectable activity.

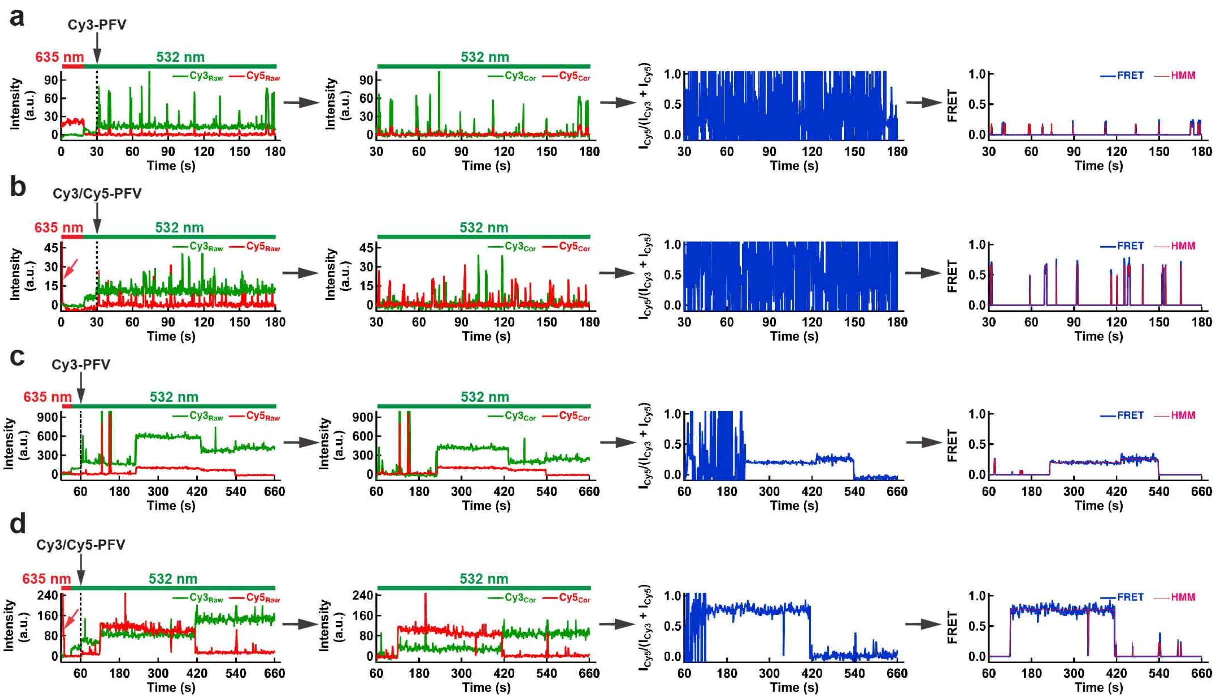

**Extended Data Figure 2. The Algorithm used to Truncate, Background Correct, and Perform HMM Analysis of the smFRET Trajectories.** (**a**) A representative TCC trajectory for Cy3-PFV collected at 100 ms frame rate. The raw intensity trajectory shows the initial 635 nm laser excitation, switching to the 532 nm laser, and real-time intasome injection (Panel 1). The beginning part of the raw intensity trajectory was truncated at the intasome injection, and Cy3, Cy5 emission backgrounds were corrected (Panel 2). The ratiometric FRET for the truncated trajectory shows erratic fluctuations when both Cy3 and Cy5 signals approach zero (Panel 3). Therefore, zero values were assigned for calculated FRET and the HMM fitting when Cy3 or Cy5 intensity reaches the background noise level (Panel 4). (**b**) A representative TCC trajectory for Cy3/Cy5-PFV collected at 100 ms frame rate. Similar analysis was performed as described in (**a**), except the initial 635 laser exposure was used to photobleach the Cy5 emission on the substrate (red arrow in Panel 1). (**c**) A representative STC trajectory for Cy3-PFV collected at 1 s frame rate. Similar analysis was performed as described in (**a**). (**d**) A representative STC trajectory for Cy3/Cy5-PFV collected at 1 s frame rate. Similar analysis was performed as described in (**a**), except the initial 635 nm laser exposure was used to photobleach Cy5 on the substrate (red arrow in Panel 1).

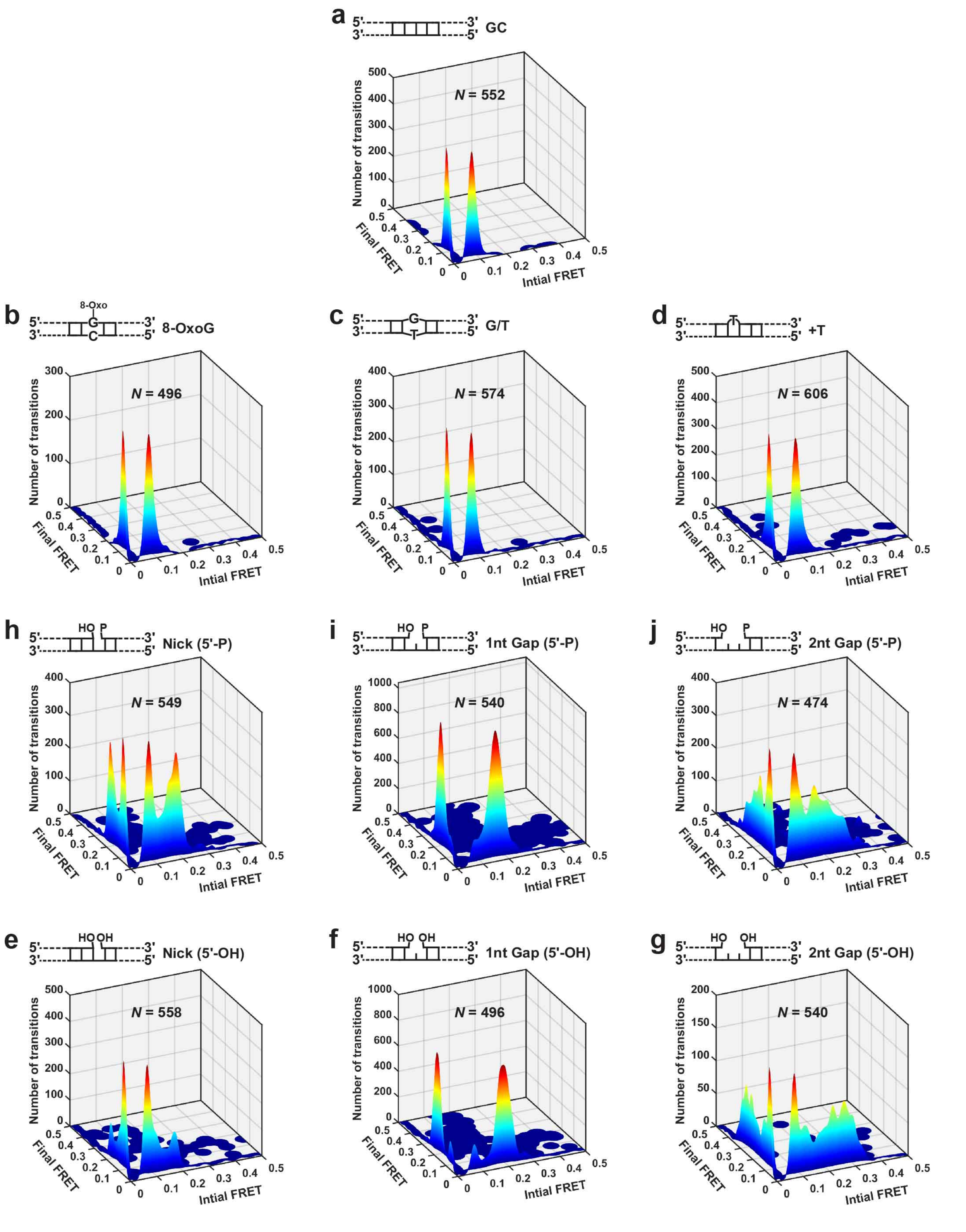

**Extended Data Figure 3. Transition Density Plots for Cy3-PFV TCC Formation with Different DNA Targets.** (**a-g**) **Transition density plots (**TDP) show the number of transitions between a given initial and a final FRET state. The identities of DNAs and the number of DNA molecules (*N*) used to build each TDP are shown.

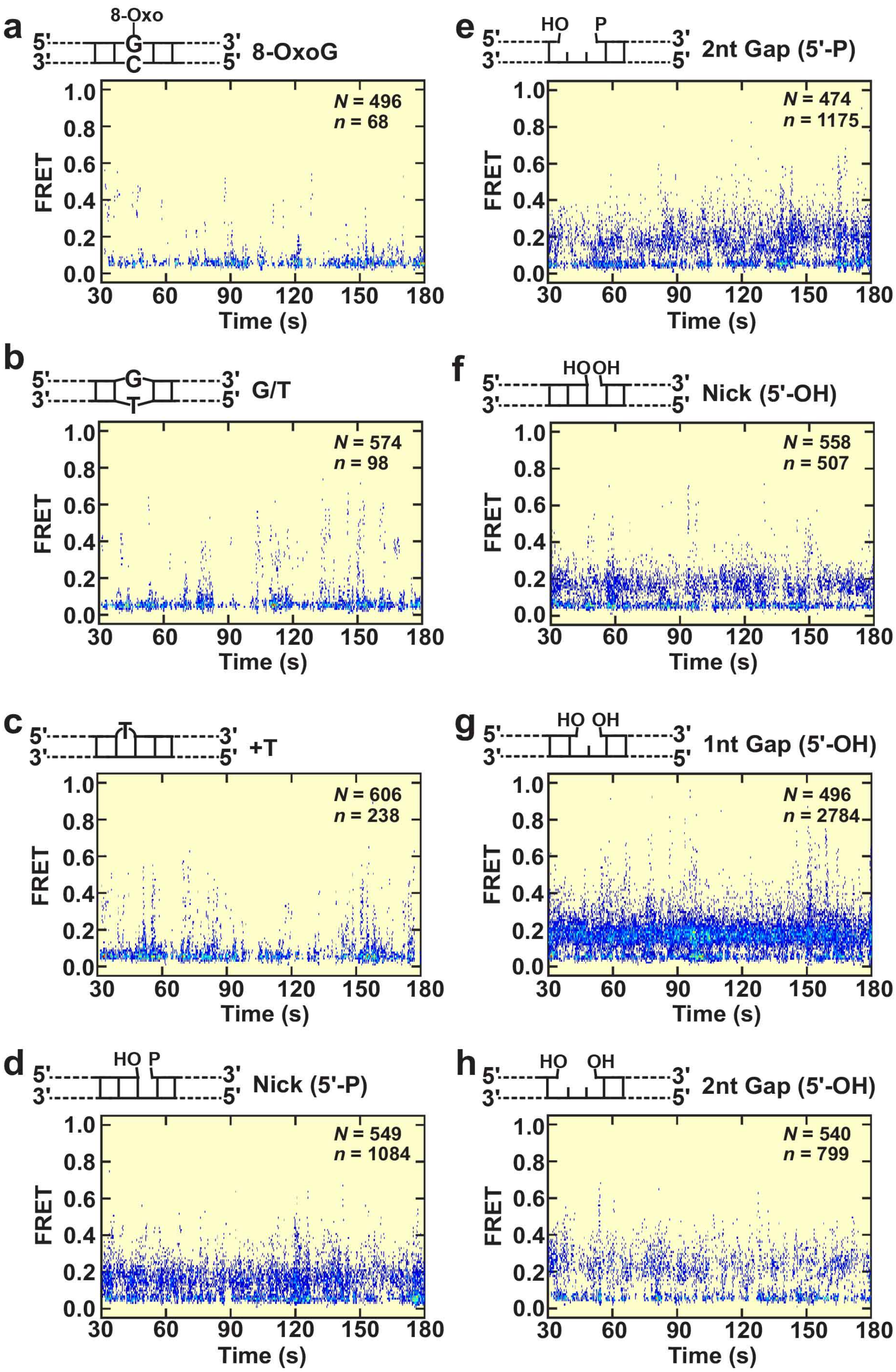

**Extended Data Figure 4. Post-Syncronized Histograms of smFRET Trajectories Showing Cy3-PFV TCC Formation on Different DNA Targets.** (**a-h**) Post-syncronized histograms (PSH) were generated by aligning smFRET trajectories from several (*N*) DNA molecules. The identities of DNAs and the total number of transitions (*n*) that crossed >0.15 FRET threshold is also shown.

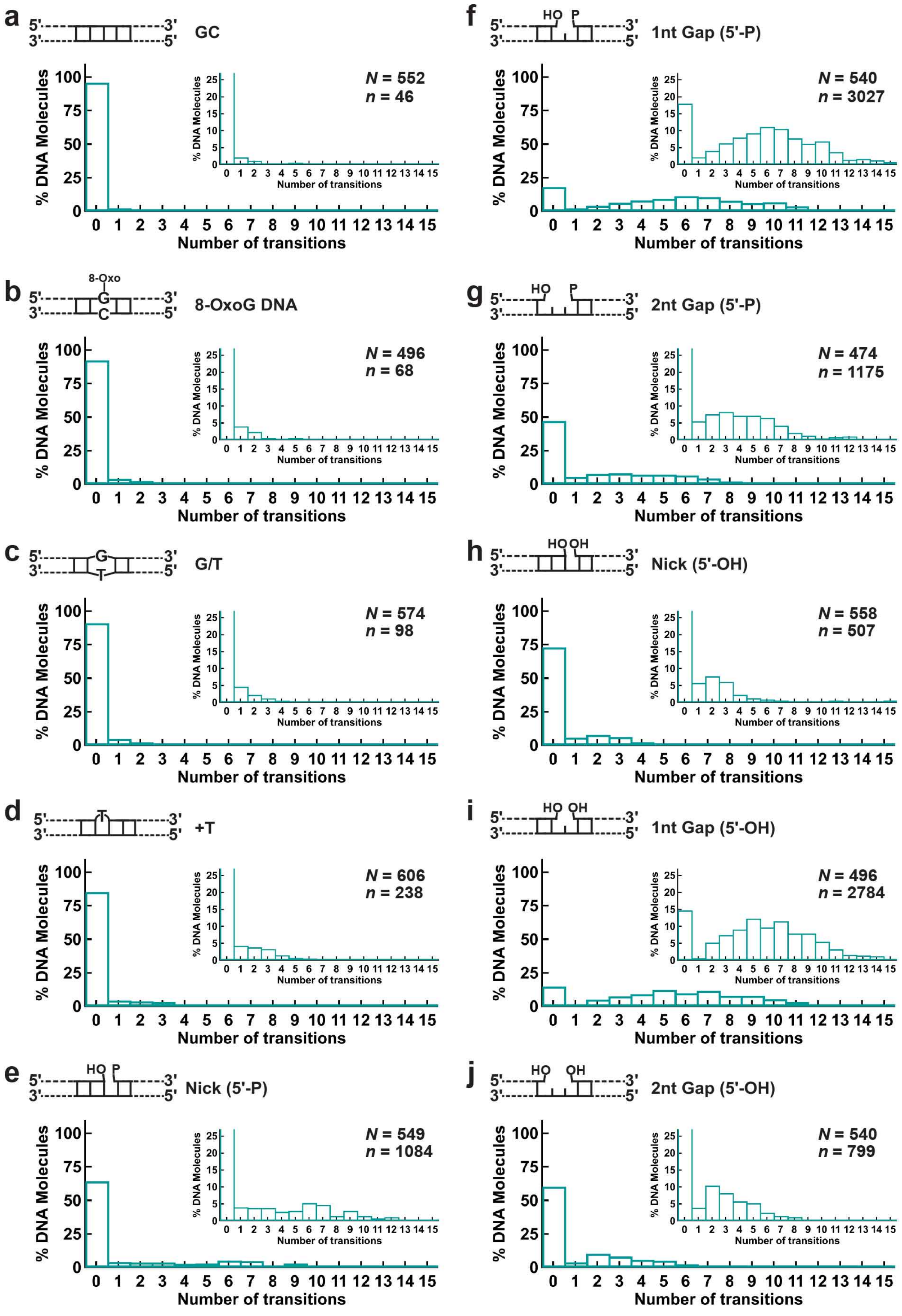

**Extended Data Figure 5. Transition Count Histograms of TCC trajectories collected at 100 ms resolution.** (**a-j**) Transition count histograms (TCH) depict the fraction of DNA molecules that show a determined number of transitions during an observation window of 2.5 min. The identities of DNAs and the total number of DNA molecules (*N*) are shown.

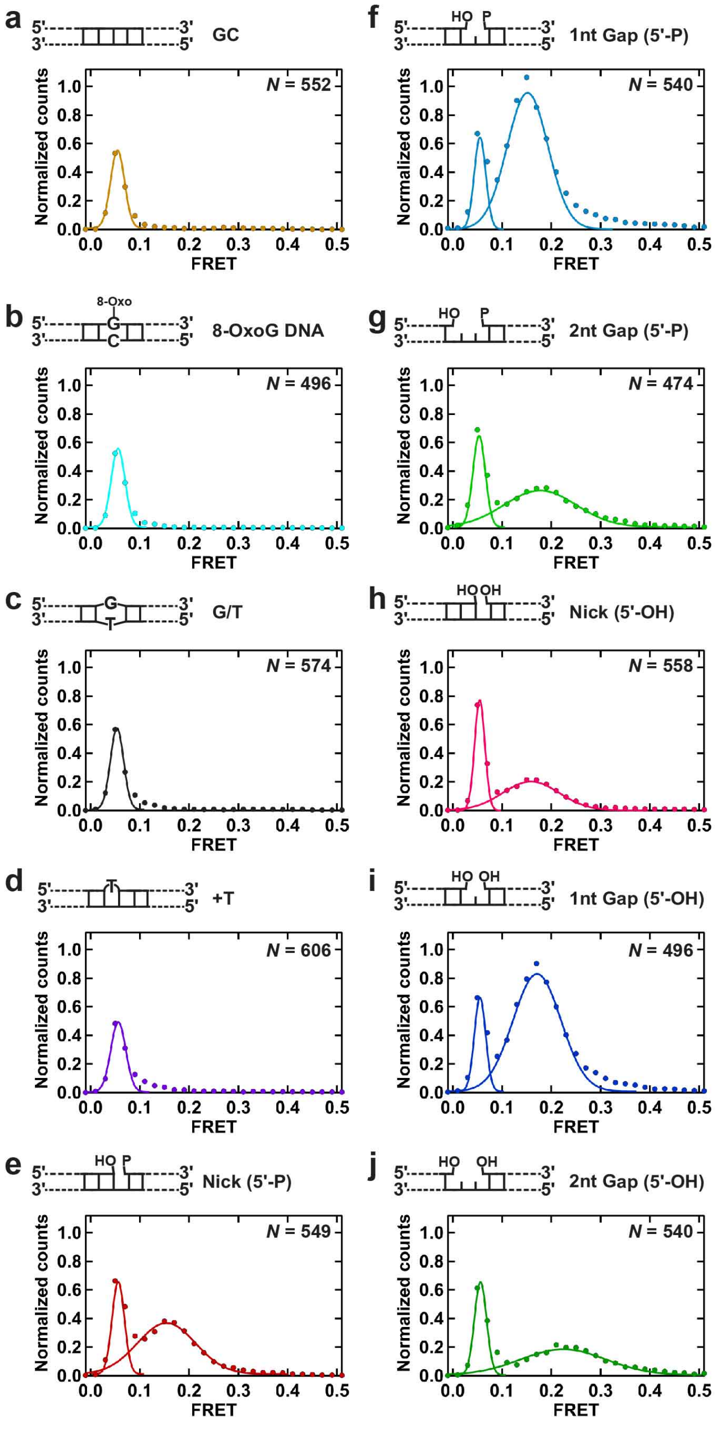

**Extended Data Figure 6. Gaussian Fittings of the TCC Histograms Collected at 100 ms Frame Rate**. (**a-j**) The histograms were normalized with respect to the area under the pseudo (0.06) FRET peak as described in Online Methods. The normalized histograms for GC, 8-OxoG, G/T, +T DNAs fit well with a single Gaussian. A combination of two Gaussians adequately models the histograms for the Nick, 1 nt and 2 nt Gap substrate, where the lower FRET peak corresponds to pseudo-FRET, and the higher FRET peak corresponds to intasome stalling on DNA near the lesion. The number of DNA molecules (*N*) included in each histogram.

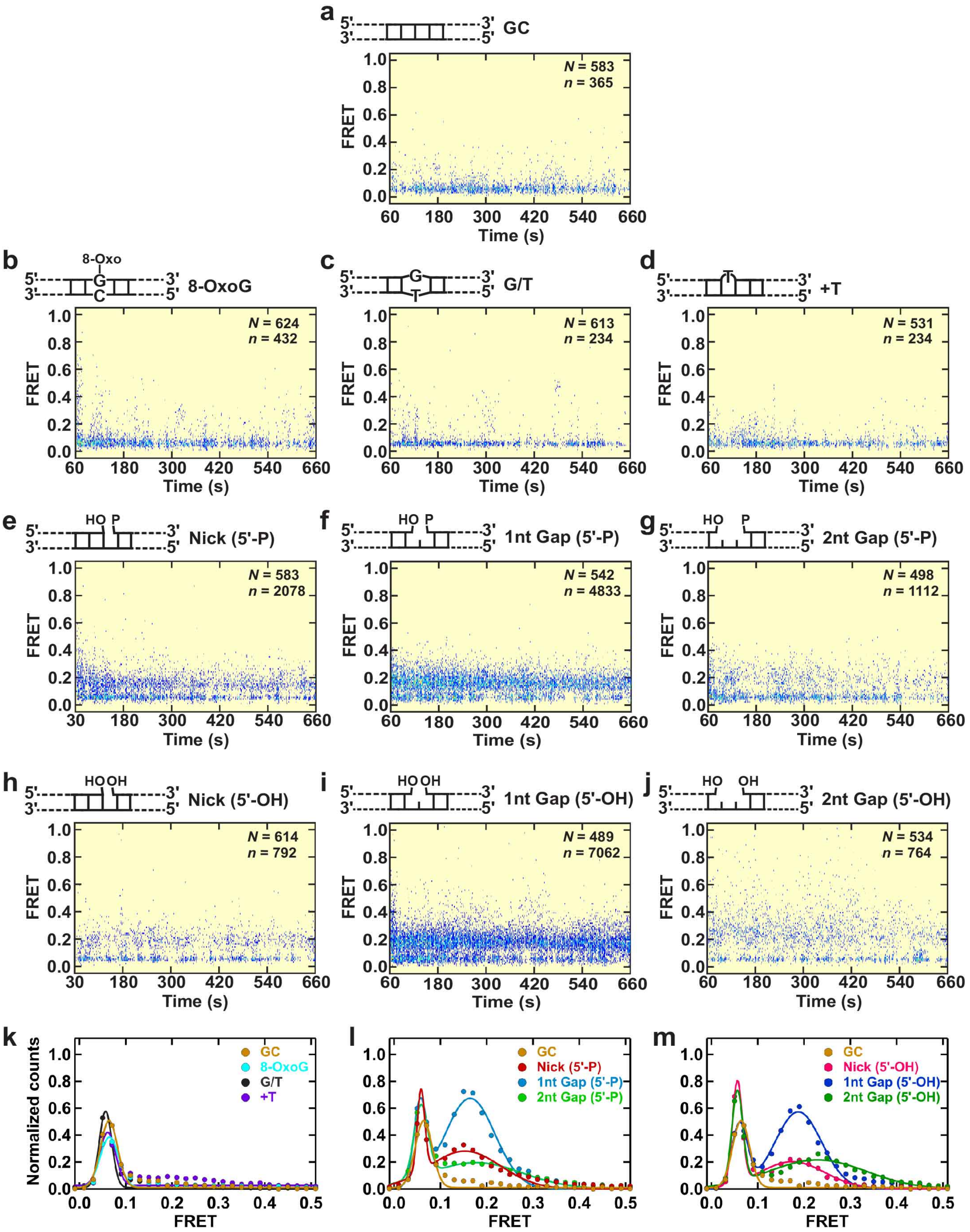

**Extended Data Figure 7. PFV Intasome TCC Dynamics at 1 s Frame Rate.** (**a-j**) Post-synchronized histograms displaying intasome interactions with different target DNA substrates. *N* is the number of DNA molecules analyzed for each substrate. *n* is the total number of transitions that crossed >0.1 FRET threshold. (**k**) Normalized smFRET histograms and their Gaussian fits showing the distributions of $E_{pseudo}$ for GC, 8-OxoG, G/T, +T DNAs. (**l,m**) Normalized smFRET histograms and their Gaussian fits showing the distributions of $E_{pseudo}$ and $E_{TCC}$ for (5'-P) (**l**) or (5'-OH) (**m**) Nick and Gap target DNA substrates.

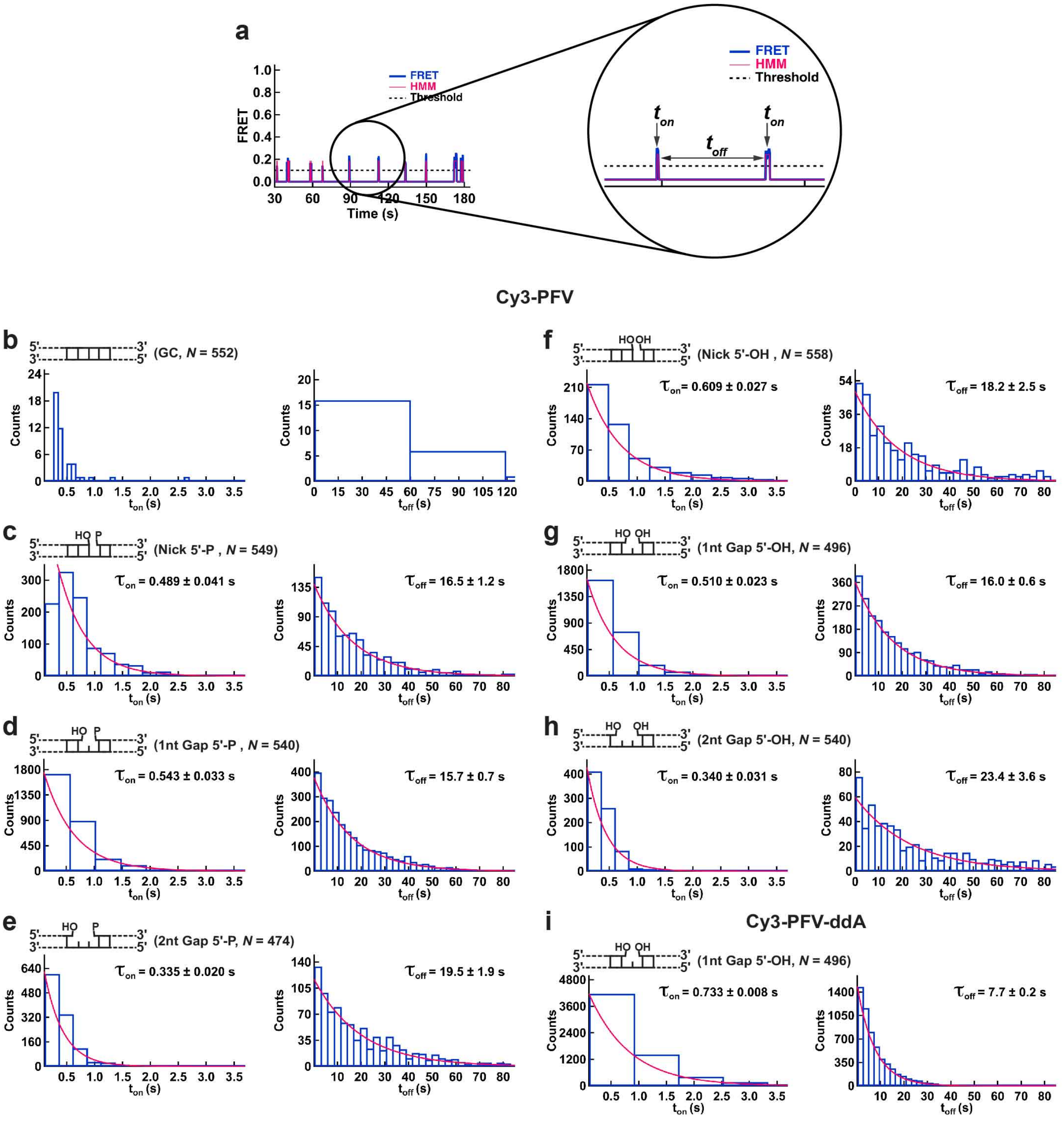

**Extended Data Figure 8. Dwell-Time Analysis for PFV Intasome TCC at 100 ms Frame Rate.** (**a**) An illustration of FRET trajectories showing the transient TCC by Cy3-PFV. The magnified view illustrates the time spent in the bound ($t_{on}$) and the unbound ($t_{off}$) states. (**b-h**) Distributions of $t_{on}$ and $t_{off}$ for different target DNA substrates. (**i**) Distributions of $t_{on}$ and $t_{off}$ for Cy3-PFV-ddA on 1nt Gap (5'-OH) DNA. The single exponential fits (red lines), the resulting average dwell times, and the errors from the fittings are shown for each distribution. The identities of the DNAs and the number of molecules ($N$) used to build each histogram are also shown.

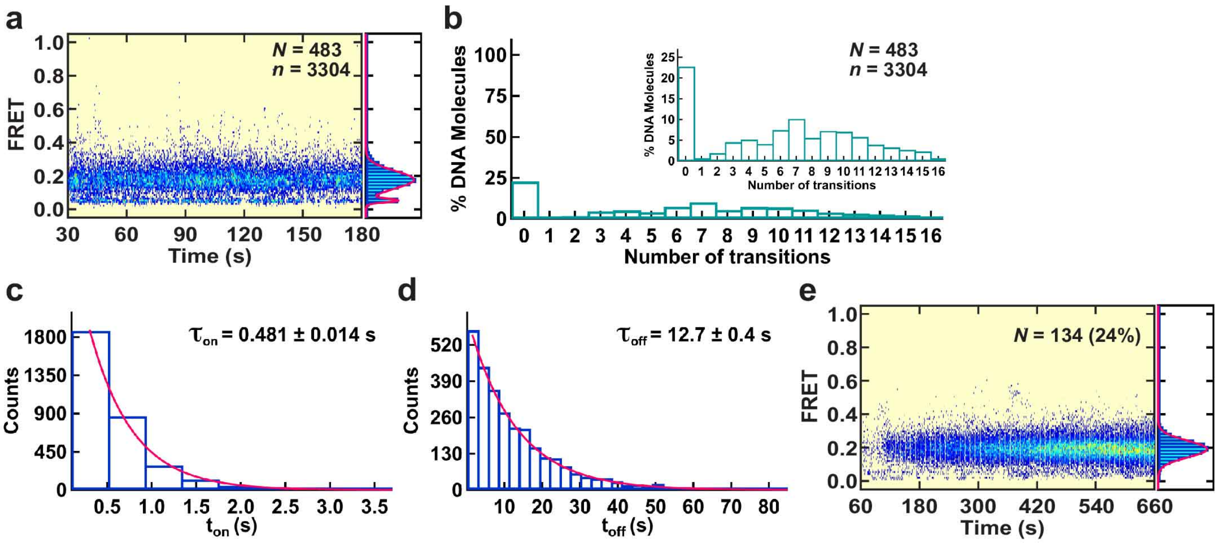

**Extended Data Figure 9. Interactions of Cy3-PFV with Blocked End Target DNA.** (**a**) The post-synchronized histograms (PSH) and the smFRET histogram corresponding to a collective of TCC events for blocked-end 1nt Gap (5'-OH) DNA. The Gaussian fits to the histograms is shown as a red line. Transition count histogram showing the fraction of DNA molecules that showed a given number of transitions during the observation window of 2.5. (**c,d**) Distributions of $t_{on}$ and $t_{off}$ for the TCC events. The single exponential fits (red lines), the resulting average dwell times, and the errors from the fittings are shown for each distribution. (**e**) The PSH and the smFRET histogram corresponding to *n* = 131 strand transfer events into blocked end 1nt Gap (5'-OH) DNA recorded at 1 s frame rate. The frequency of strand transfer (%) is shown in parenthesis.

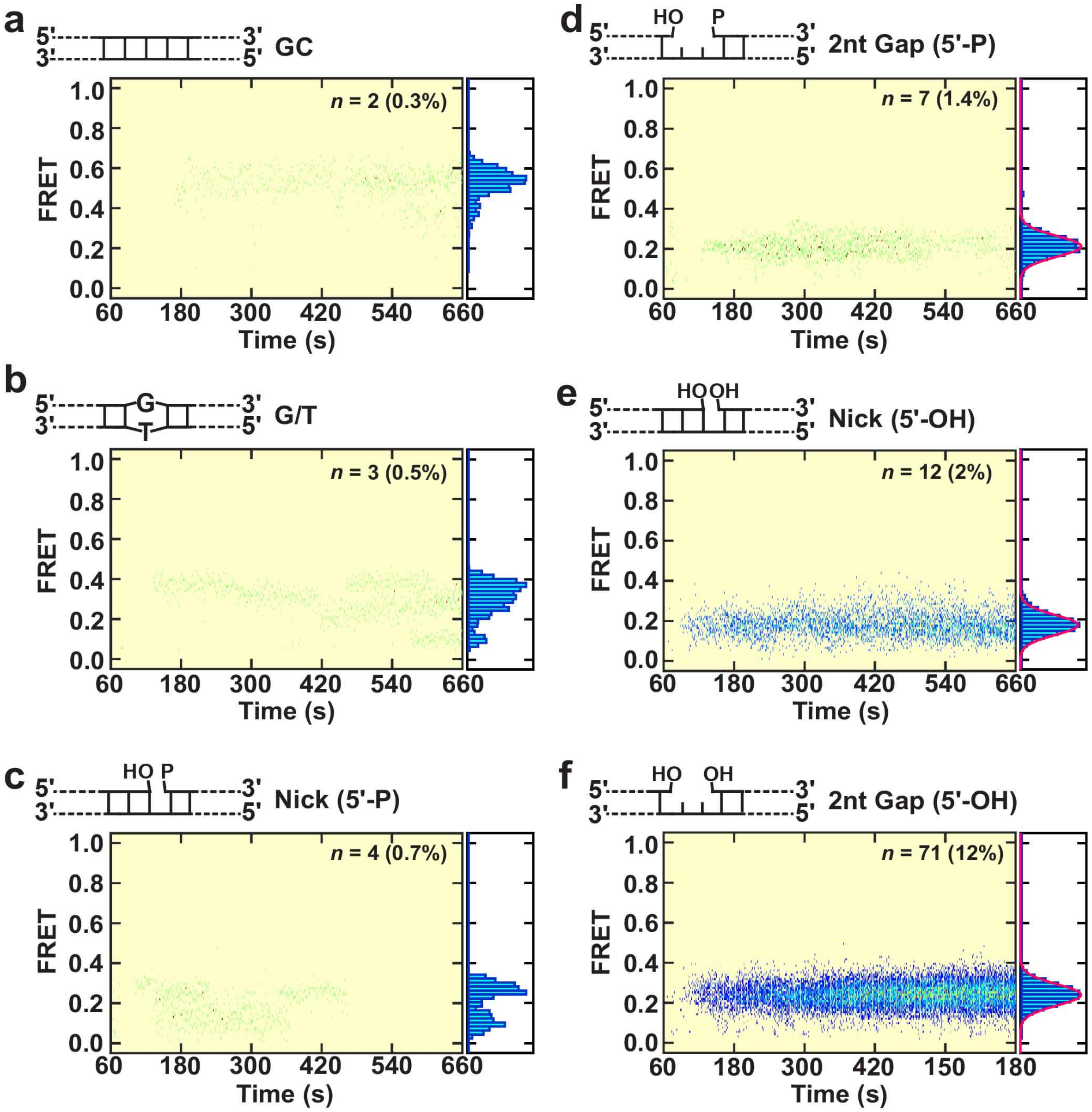

**Extended Data Figure 10. Post-Syncronized Histograms and smFRET Histograms of Cy3-PFV Strand Transfer Events.** (**a-f**) The post-synchronized hostogrames (PSH) were generated by aligning smFRET trajectories that showed the formation of a strand transfer complex (STC). The identities of DNAs, the total number of strand transfer events (*n*) included in each analysis, and the efficiencies of STC formation (%) are shown. The Gaussian fits to the smFRET histograms are shown as red lines.

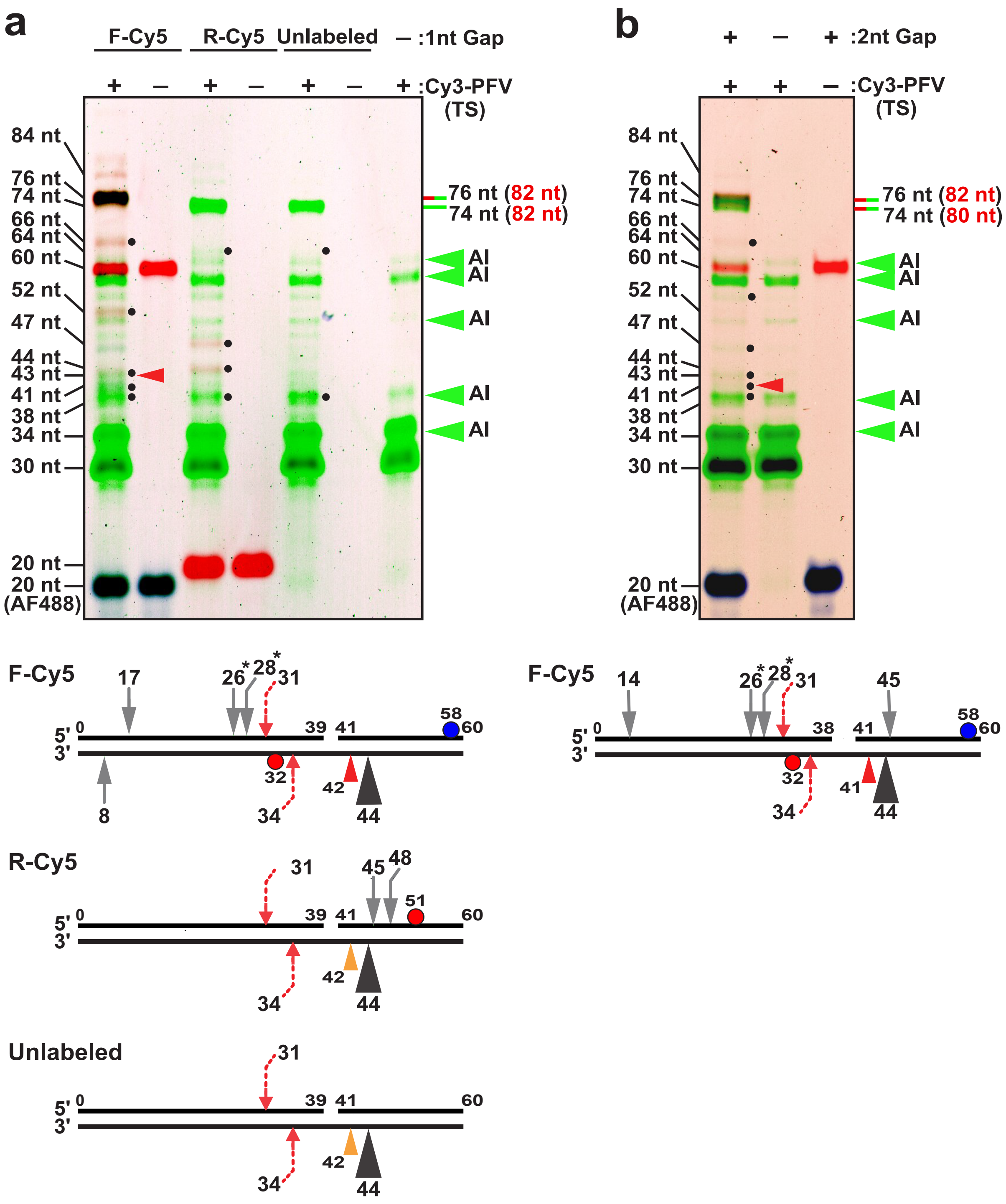

**Extended Data Figure 11. Enhance Image Contrast of Cy3-PFV (TS) Integration into 1 nt Gap (5'-OH) and 2 nt Gap (5'-OH) Target DNA. a)** (top) Enhance image contrast of 1 nt Gap (5'-OH) denaturing PAGE gel shown in Fig. 3d. Major half-site integration product of 74 nt and 76 nt (with Cy5 fluorophore that increases apparent size by 2 nt) are marked with green and green+red lines, which are equivalent to 82 nt integration product of unlabeled PFV intasome with the F-Cy5 target DNA shown in Fig. 3 b,c. Dots next to gel band indicate integration products mapped to the DNA substrates illustrated below following subtraction of 30 nt corresponding to the ligated vDNA. Red arrowhead marks the 42 bp aborted strand transfer product (see Fig. 3 b,c) on both the gel above and integration site illustration below (orange arrowhead in illustration shows the location of undetectable aborted integration product since the corresponding DNA strand does not contain a fluorophore label). AI indicates auto-integration products found in both control and target DNA lanes. (bottom) Illustration F0Cy5, R-Cy5 and unlabeled target DNAs with calculated integration sites as arrowheads or arrows. The major half-site product maps to 44 nt on the non-lesion containing strand (black arrowhead), while the corresponding concerted integration event would occur at 40 nt on the lesion-containing strand where a phosphate bond required for the isoenergetic strand transfer is absent. Gray arrows in bottom illustration indicate the location of half-site products (* indicate products most likely location but with a second possible calculated location at 11 and 13 nt on the same strand). Red dashed arrows show the location of the most frequent concerted integration product that appears with all three target DNAs. The 44 nt half-site event accounts for >90% of the products, while all other mapped events account for <5% of the total products. (**b**) Enhanced contrast of denaturing PAGE gel of Cy3-PFV integration products into F-Cy5 containing a 2 nt Gap (5'-OH). Markings of gel above and illustration below are similar to Panel a except with reference to 2 nt Gap shown in Fig. 3 b,c. As will Panel a, the 44 bp half-site events accounts for >90% of the integration products.

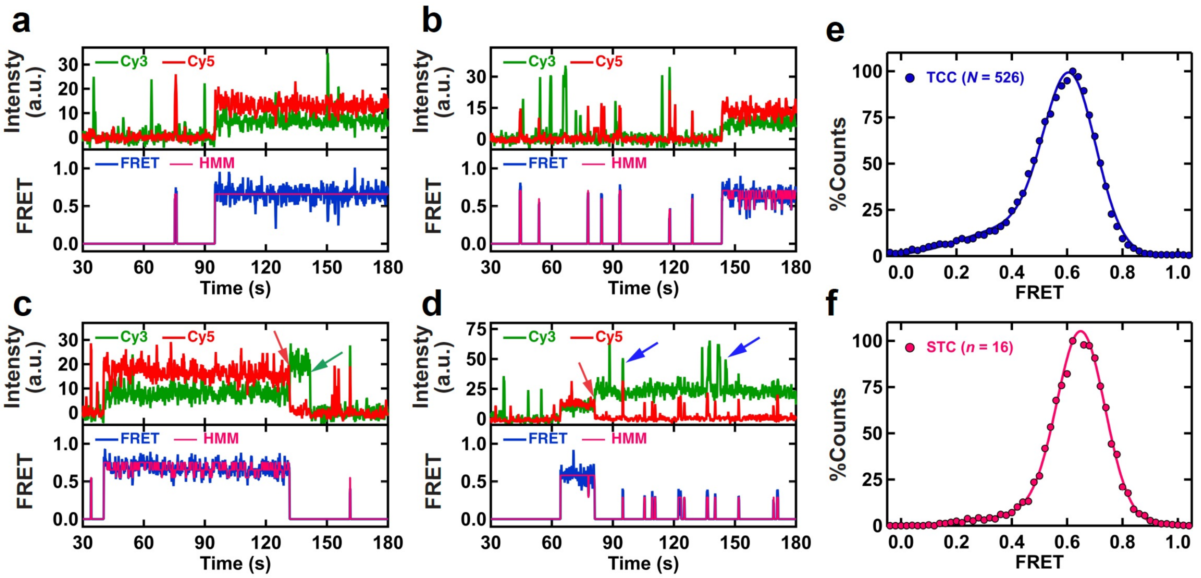

**Extended Data Figure 12. Probing the structural dynamics of PFV intasomes during target capture and strand transfer.** (**a-d**) Representative intensity trajectories and the resultant FRET trajectories with the HMM fits showing strand transfer by Cy3/Cy5-PFV into 1nt Gap (5'-OH) DNA. For the longer STC FRET states, Hidden Markov Model (HMM) analysis often fits fluctuations that are not originating from anti-correlated Cy3, Cy5 intensities. The photobleaching of Cy3 and Cy5 are denoted by green and red color arrows, respectively. The blue color arrows indicate aggregate excursions of the intasome that occasionally produce pseudo-FRET transitions. (**e**) smFRET histogram with the Gaussian fit for the TCC events exhibited by Cy3/Cy5-PFV binding to 1nt Gap (5'-OH) DNA. (**f**) smFRET histogram with the Gaussian fit for STC events exhibited by Cy3/Cy5-PFV into 1nt Gap (5'-OH) DNA. Data was collected at 100 ms frame rate. The total number of DNA molecules (*N*) examined or the total number of STC events (*n*) are included in each histogram is indicated.
